# Evolutionary invasion analysis for structured populations: a synthesis

**DOI:** 10.64898/2026.03.23.713828

**Authors:** Ryosuke Iritani, Troy Day

## Abstract

Natural populations exhibit complex class structures that fundamentally shape evolutionary dynamics. Yet predicting adaptation in such systems remains a central challenge in ecology because ecological realism inherently generates high-dimensional models that obscure biological interpretation and hinder general evolutionary prediction. Here, we overcome this long-standing obstacle by developing structural evolutionary invasion analysis, a unifying theoretical framework integrating invasion determinants with the Projected Next-Generation Matrix (PNGM). The invasion determinant provides a general, closed-form algebraic condition for invasion, while the PNGM structurally compresses life-cycle graphs by eliminating non-focal classes. Crucially, this compression formally derives timescale separation, rigorously preserves Fisher’s reproductive values, and guarantees that the location, convergence, and evolutionary stability of equilibria remain identical to those of the full ecological system. By applying the framework to examples spanning evolutionary epidemiology, stage structure, and inclusive fitness, we show how to capture the direct impact of ecological complexity on evolutionary outcomes while retaining analytical tractability. Our results establish a general principle linking ecological structure to evolutionary prediction, enabling ecologists to analyze adaptation and eco-evolutionary dynamics in heterogeneous populations.

## 1 Introduction

Natural populations exhibit complex class structures (e.g., sex, age, body size, or habitat) that fundamentally shape their ecology, demography and life histories. Theoretical models reveal that state-dependent selection drives diverse eco-evolutionary phenomena, from dispersal and social behaviours to pathogen defences (e.g., Rodrigues & Gardner 2013; Iritani & Iwasa 2014; Rodrigues & Gardner 2015; Úbeda & Jansen 2016; Iritani & Cheptou 2017; Boots & Best 2018; Iritani *et al*. 2019; Kuijper & Johnstone 2019; Iritani 2020; Iritani *et al*. 2021; Abe *et al*. 2021; Buckingham *et al*. 2023). Empirical research corroborates these predictions, quantifying how the patterns of natural selection vary across classes and ecological contexts (e.g., Coulson *et al*. 2005; Pelletier *et al*. 2007; Kruuk & Hill 2008; Smallegange & Coulson 2013). Because state-dependent selection directly feeds back into population growth and stability, elucidating how class structure modulates natural selection is fundamental for predicting both evolutionary and ecological dynamics in realistic biological systems.

To analyze state-dependent selection under complex ecological heterogeneity, researchers synthesize adaptive dynamics and matrix population models (Metz *et al*. 1992; Takada & Nakajima 1992; Dieckmann & Law 1996; Geritz *et al*. 1998; Caswell 2001; Rees & Ellner 2016; Williams & Kamel 2021). This “evolutionary demography” framework translates demographic processes into the invasion condition of a rare mutant lineage in ecological environments set by a resident lineage. Crucially, invasion depends not merely on immediate reproduction, but on long-term demographic contributions quantified as Fisher’s (1930) reproductive values (Fisher 1930; Taylor 1990; Caswell 2001; Rousset 2004; Lion 2018; Lion & Gandon 2022). However, while global databases (e.g., `COMPADRE` and `COMADRE`) provide rich empirical data (Salguero-Gómez *et al*. 2014, 2016; *Jones et al*. 2022), the high dimensionality of these matrices hinders transparent analytical insights. Consequently, studies frequently rely on numerical approaches rather than explicit analytical solutions (e.g., Coulson *et al*. 2010, 2011; Smallegange & Coulson 2013). Although powerful for specific scenarios, these numerical approaches obscure underlying life-history trade-offs and limit broad mechanistic generalizations.

To derive analytically tractable and biologically interpretable invasion conditions from these demographic models, Hurford *et al*. (2010) adapted the next-generation matrix (NGM) approach from mathematical epidemiology (Diekmann *et al*. 1990; van den Driessche & Watmough 2002). By transforming the Jacobian to capture lifetime reproductive output, the NGM yields a systematic invasion condition based on the basic reproduction number, *w* > 1. Evaluating this threshold often provides deeper biological insights into long-term evolutionary consequences.

Despite its utility, the NGM approach often yields invasion conditions with obscure biological interpretations, particularly in high-dimensional systems. For instance, even in a simple two-class model, the invasion condition emerges as an opaque quadratic equation (e.g., Massol & Cheptou 2011; Massol & Débarre 2015; Zurita-Gutiérrez & Lion 2015). Naturally, as the number of population classes increases, obtaining simple, analytical expressions becomes increasingly elusive.

To mitigate this dimensionality problem, various reduction methods have been developed, though primarily within ecological or epidemiological contexts (Caswell 2001; de Camino Beck & Lewis 2007; Rueffler & Metz 2013; Lewis *et al*. 2019). While some techniques have been introduced to evolutionary questions such as linear-algebraic methods (Rueffler *et al*. 2012) or the type-reproduction number (Roberts & Heesterbeek 2003), they do not provide a systematic framework to derive biologically interpretable invasion conditions. Consequently, the application of dimension reduction to evolutionary invasion analysis remains underdeveloped.

In this article, we resolve a pervasive limitation in eco-evolutionary theory by formulating structural evolutionary invasion analysis, a general framework for analysing adaptation in class-structured populations. By integrating invasion determinants with the Projected Next-Generation Matrix (PNGM), our approach links ecological structure directly to evolutionary invasion criteria. Evaluated through Fisher’s (1930) reproductive value (Taylor 1990; Caswell 2001; Rousset 2004) and interpreted using life-cycle graphs (Caswell 2001; Rueffler *et al*. 2012; Giaimo *et al*. 2018), the framework reveals how complex demographic processes can be reduced without altering evolutionary outcomes. Applied across four illustrative examples under weak selection, our synthesis demonstrates how analytically intractable eco-evolutionary models become interpretable predictors of adaptation. Rather than providing a new modelling tool alone, structural evolutionary invasion analysis establishes a general principle for connecting ecological complexity with evolutionary prediction.

## 2 Overview: Evolutionary invasion analysis

We here review the next-generation matrix (NGM) methodology for evolutionary invasion analysis (Hofbauer & Sigmund 1990; Dieckmann & Law 1996; Geritz *et al*. 1998; Hurford *et al*. 2010). We focus on continuous-time models of class-structured populations and present discrete-time version in Examples (Table 1).

**Table 1:**
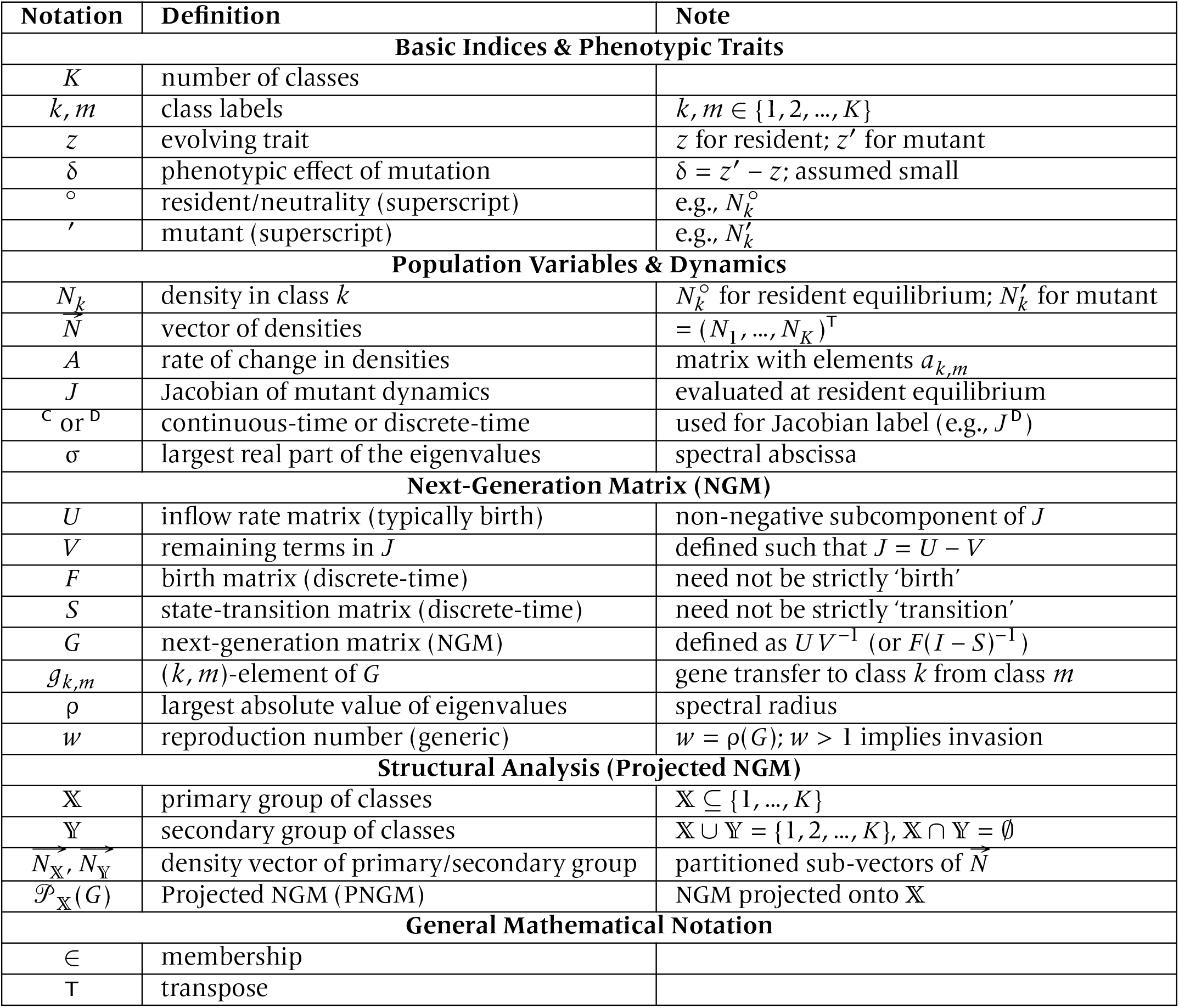
Summary of notation used in the main text.

Following standard adaptive dynamics, we assume a separation of ecological and evolutionary timescales. A rare mutant with a slightly deviated phenotype *z*′ = *z* + δ (weak selection) is introduced into a monomorphic resident population at equilibrium with trait *z*. Because “invasion implies substitution” under these conditions (Geritz 2005; Priklopil & Lehmann 2020), the long-term sequence of evolutionary substitutions can be fully characterized by evaluating the mutant invasion condition.

### Ecological dynamics

Consider a population structured into *K* discrete classes (e.g., habitats, stages, sizes, or epidemiological states). For any pair of classes *k* and *m*, we assume that an individual of class *m* can produce class *k*-individuals, either directly (via reproduction or transition) or indirectly (via other classes). Such class structure is called irreducible (Caswell 2001; Stott *et al*. 2010, Figure 1A).

**Figure 1:**
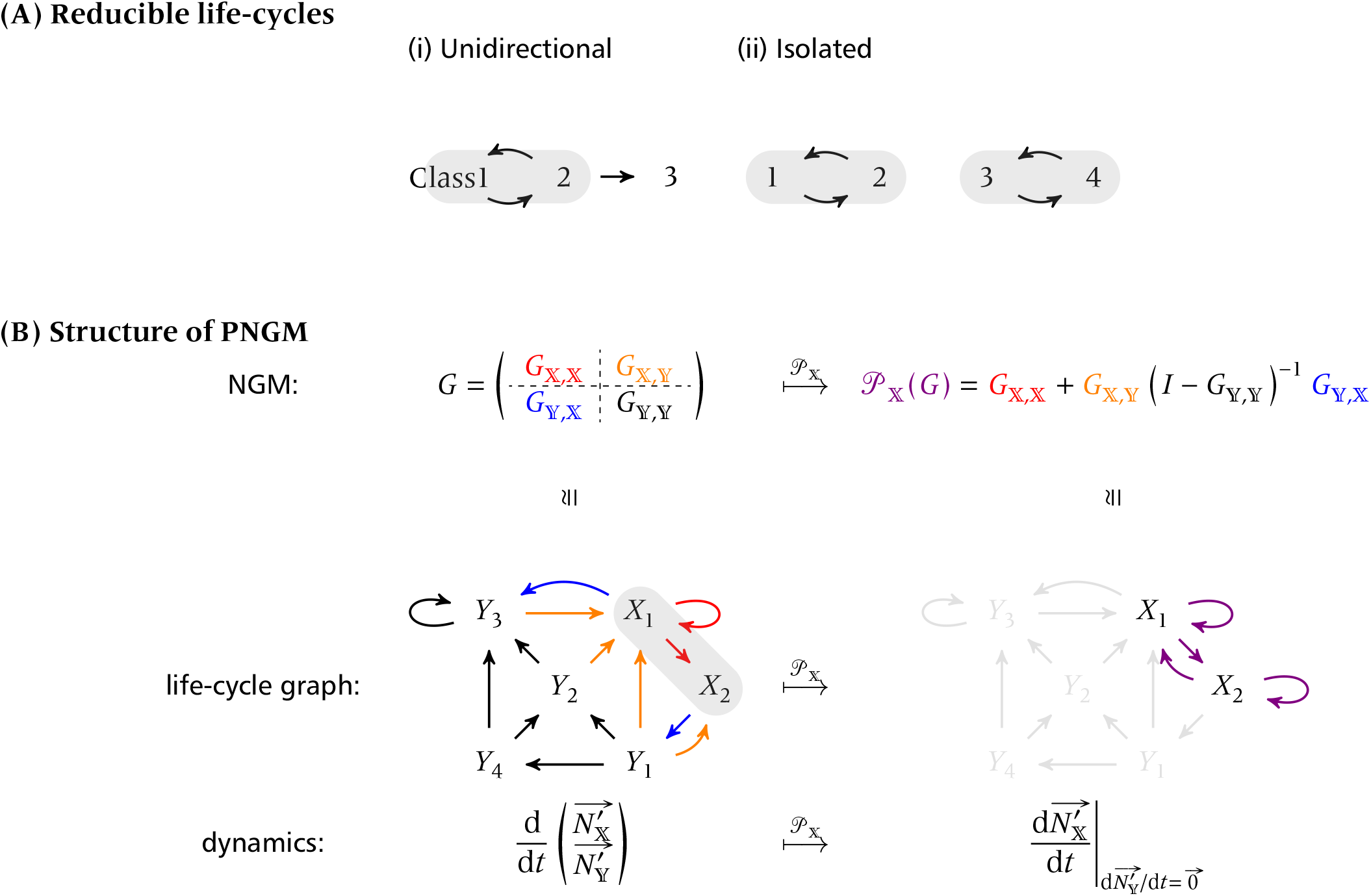
A schematic illustration of (A) reducibility and (B) PNGM. (A) Two typical examples of reducible life-cycle graphs (i.e., non-irreducible life-cycles). Arrows represent transitions between classes, and gray bands highlight subsets of classes that form irreducible components — subgraphs in which every node is reachable from any other within the same component. In both examples, the invasion-implies-substitution principle fails: a mutation arising in class 3, for instance, cannot spread to fix in the entire population in both panels: (i) Classes 1 and 2 are interconnected, but no gene flow from class 3 to class 2 is possible. Thus a mutant emerging in class 3 will never get fixated. (ii) Classes 1 and 2 form one interconnected group, and classes 3 and 4 form another. There is no gene flow between these two groups, resulting in two isolated populations, and thus fixation is impossible. (B) We write 𝕏 = {*X*_1_, *X*_2_} for primary and 𝕐 = {*Y*_1_, *Y*_2_, *Y*_3_, *Y*_4_} for secondary groups. The colors: red for the pathway [𝕏 ← 𝕏], orange for [𝕏 ← 𝕐], blue for [𝕐 ← 𝕏], and black for [𝕐 ← 𝕐]. Left panels are the original structures (NGM, life-cycle graph, and mutant dynamics from top to bottom), and right panels are projected structures.

We visualize model structures using a life-cycle graph (Caswell 2001), where vertices represent classes and directed edges (paths) represent lineage propagation via class-transition or births. The irreducibility assumption then means that for any pair of classes, (*k, m*), at least one finite-length path exists from the vertices *k* to *m* (‘strongly connected’ life-cycle graph).

The rate of change in class densities is modelled using ordinary differential equations (ODEs):

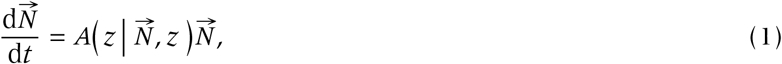

where 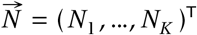 collects the class densities and *A* represents a matrix, as a function of the resident trait and densities, of the rates of density changes. We assume that, if the population is monomorphic with a resident trait value *z*, then there exists a unique, globally stable ecological equilibrium 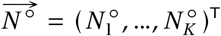 with 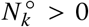 for all classes, *k* = 1, 2, …, *K* and 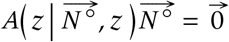 (zero vector). We refer to this equilibrium as the resident equilibrium.

Note that the potential existence of alternative equilibria or attractors does not affect our analysis, since the weak selection assumption prevents the population from converging to an alternative attractor (see Geritz *et al*. 2002; Priklopil & Lehmann 2020). We thus assume that the resident population dynamics are regulated by sufficient negative density-dependence at high densities, and that the population can grow at low densities. Under these standard ecological conditions, the resident dynamics admit a strictly positive, globally stable equilibrium (Cushing 1998).

### Dynamics of a rare mutant

Suppose a mutation arises with phenotype *z*′ = *z* + δ (where |δ| ≪1). The mutant density in class *k*, 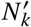, changes according to:

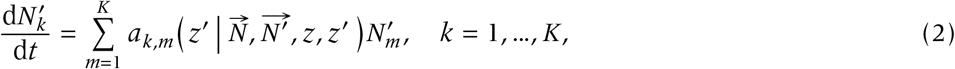

where *a*_*k,m*_ is the per-capita rate at which class-*m* mutants produce to class-*k* mutants. Assuming the mutant is initially rare, we linearize the system around the resident equilibrium 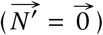:

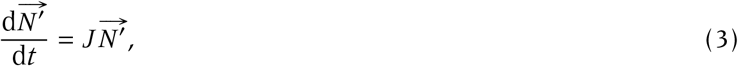

where the Jacobian *J* has entries 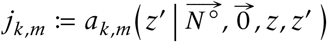. The mutant successfully invades if the spectral abscissa (the largest real part of the eigenvalues) σ(*J*) is positive.

Directly computing the spectral abscissa, σ(*J*), is typically intractable for systems with more than two classes. Furthermore, ad-hoc algebraic derivations (such as evaluating the determinant of *J*) often yield model-specific conditions devoid of biological transparency, driving the need for more systematic approaches.

Most alternatives to analyzing σ(*J*) involve shifting the timescale of evolutionary dynamics. Rather than tracking the mutant population over absolute time (e.g., days), these approaches evaluate population change over an organismal generation. This concept is best illustrated using a simple unstructured population. Suppose the mutant density *n*′ changes according to the scalar ODE d*n*′/d*t* = *un*′ − *vn*′, where *u* and *v* are the constant per-capita birth and death rates, respectively, and time is measured in days. The spectral abscissa is simply σ = *u* − *v*, which has units of days^−1^. Invasion occurs if *u* − *v* > 0, meaning the daily fecundity exceeds the daily mortality. However, this inequality can be algebraically rearranged as *uv*^−1^ > 1. The left-hand side is now a dimensionless quantity, rendering it independent of the absolute time units. Biologically, because a mutant individual experiences a constant mortality rate *v*, its expected lifespan is *v*^−1^ days. Multiplying this expected lifespan by the daily birth rate, *u*, yields the dimensionless ratio *w* := *uv*^−1^, which represents the expected number of offspring a mutant individual will produce over its lifetime. If this lifetime reproductive output exceeds unity, the mutant population will grow and successfully invade.

While this unstructured example is simple enough that both invasion criteria (σ(*J*) > 0 and *uv*^−1^ > 1) can be readily computed and biologically interpreted, introducing class structure significantly complicates the analysis. In class-structured systems, the transition from continuous-time rates to generational reproductive output is less straightforward, necessitating the formal mathematical machinery of the next-generation matrix.

### Next-generation matrix (NGM)

The NGM method generalizes the concept of per capita lifetime reproductive output to structured populations. This is achieved by constructing a matrix, NGM, *G*, whose entries *g*_*k,m*_ represent the expected number of class-*k* offspring produced by a single class-*m* individual over its lifetime. To construct this matrix, we partition the continuous-time Jacobian as *J* = *U* − *V*. Here, *U* is a non-negative matrix capturing the inflow of new individuals (e.g., “births”) into each class, while *V* captures all mortality and transition rates among the classes. From a mathematical standpoint, the biological labels assigned to these transitions are somewhat arbitrary, meaning the partitioning *J* = *U* − *V* is not necessarily unique. Although we will explore the implications of this non-uniqueness later, any valid partition must satisfy three requirements: (i) *U* ≥ 0 (non-negative inflows); (ii) σ(−*V*) < 0 (no inflows *U* = *O* implies the population asymptotically approaches extinction); and (iii) *V* ^−1^ exists and is non-negative.

Under these conditions, the NGM is defined as *G* := *U V* ^−1^, which satisfies:

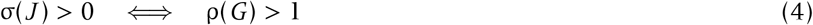

(Hurford *et al*. 2010), where ρ(*G*) denotes the spectral radius of the NGM (i.e., its largest eigenvalue in modulus), and the left equality holds if and only if so does the right. Consequently, we can evaluate the invasion condition equivalently using either σ(*J*) or ρ(*G*). One can verify that the earlier scalar ODE example represents a special, one-dimensional case of this general result.

While the NGM improves interpretability over σ(*J*), standard formulations remain analytically challenging for complex life-cycles. Alternative graph-theoretical reductions (Rueffler *et al*. 2012; Rueffler & Metz 2013) are often mathematically demanding. A more direct strategy involves strategically choosing a non-standard decomposition (*J* = *U* − *V*) to simplify the NGM. However, lacking a unifying theory, this approach has historically forced researchers to rely on ad hoc, model-specific derivations.

We synthesize these approaches into *structural evolutionary invasion analysis*, a framework integrating the invasion determinant and systematic Jacobian *J* decomposition. By exploiting the life-cycle’s algebraic architecture, it structurally characterizes evolutionary dynamics. Operating under standard assumptions (weak selection, a globally stable resident equilibrium, and an irreducible NGM), this synthesis provides a tractable toolkit for complex evolutionary demographic models.

## 3 Synthesis: structural evolutionary invasion analysis

The first component of our framework is an algebraic tool we term the “invasion determinant”, − det(*I* − *G*) (Appendix A). Although similar determinant-based conditions appear in previous works (Taylor & Bulmer 1980; Courteau & Lessard 2000; Day & Burns 2003; Metz & Leimar 2011; Rueffler *et al*. 2012), they remain widely underutilized. Evaluating this determinant directly captures the growth potential of a rare mutant lineage, yielding a concise biological threshold for invasion.

The second component is the *Projected Next-Generation Matrix* (PNGM) approach (Appendix C). Designed to systematically reduce model-dimensionality, the PNGM builds on the epidemiological concept of the type-reproduction number (Roberts & Heesterbeek 2003; Inaba & Nishiura 2008). Its mathematical foundation is a nonnegative decomposition theorem (Appendix B; Box 1), which guarantees that any biologically motivated, non-negative partitioning of the Jacobian preserves the true invasion threshold.

By exploiting this theorem, the PNGM algebraically eliminates a subset of classes to focus solely on the remainder. Biologically, this is equivalent to a timescale separation: ‘fast’ secondary classes are set to quasi-equilibrium 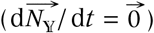 and removed from the life-cycle graph (Box 3). Crucially, this network compression preserves both the invasion threshold and Fisher’s (1930) reproductive values of the retained classes, maintaining clear biological interpretability.

A further key property is that both methods preserve the (i) position of the evolutionary equilibrium, (ii) convergence stability towards it, and (iii) evolutionary stability against invasion of alternative mutants (Appendix D). These properties provide useful tools for predicting long-term evolutionary outcomes, extending beyond the analysis of instantaneous invasion conditions.

### Key component 1: Invasion determinant

The invasion invasion condision can be expressed as:

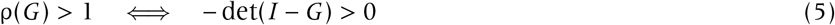

(Appendix A), where det() represents a matrix determinant. By formalizing this − det(*I* − *G*) as the “invasion determinant”, we obtain a tractable, closed-form condition for mutant invasion that is applicable to arbitrary class-structured populations.

Incidentally, we can also derive the invasion condition using the original, continuous-time Jacobian *J* :

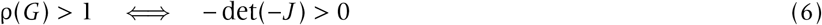

(Appendix A; Day & Burns 2003; Hastings & Botsford 2006b; Boots & Best 2018).

While the invasion determinant powerfully simplifies algebraic analysis for high-dimensional systems, it often lacks biological transparency because it obscures Fisher’s (1930) reproductive values. Nevertheless, it serves as a robust theoretical tool. Furthermore, Metz & Leimar (2011) demonstrated that this determinant condition remains valid even when relaxing the weak-selection assumption, extending its utility beyond traditional adaptive dynamics.

### Key component 2: Projected Next-Generation Matrix (PNGM)

To construct the PNGM, we first partition the classes of *G* into two mutually exclusive groups based on any relevant biological feature: a ‘primary’ group 𝕏 and a ‘secondary’ group 𝕐 (Figure 1). For example, a six-class system might be divided into 𝕏 = {*X*_1_, *X*_2_} and 𝕐 = {*Y*_1_, *Y*_2_, *Y*_3_, *Y*_4_}.

We block-partition the NGM as:

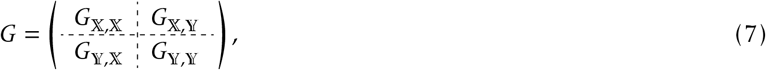

where each block represents the transfer of genes (via reproduction or transition) within and between these groups. For instance, *G*_𝕏,𝕐_ captures gene transfer from the secondary 𝕐 to the primary 𝕏 group (Figure 1).

Primary mutants produce primary offspring either directly (*G*_𝕏,𝕏_) or indirectly via the secondary group (*G*_𝕏,𝕐_ *G*_𝕐,𝕏_). Summing these pathways over *n* generations yields the expected number of primary offspring:

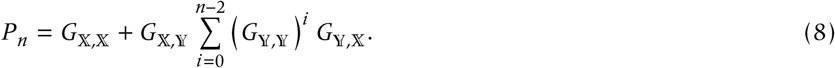

Taking the limit *n* → ∞, we define the dimensionally reduced PNGM:

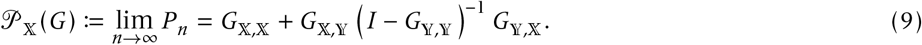

The second term effectively accumulates all indirect loops ([𝕏 ← 𝕐 ← · · · ← 𝕐 ← 𝕏]).

Notably, 𝒫_𝕏_(*G*) and *G*_𝕏,𝕏_ have the same size, and thus 𝒫_𝕏_(*G*) is the projection of *G* onto a lower dimensional matrix whose classes are given by the primary group 𝕏; also note that while ‘projection’ in population demography often refers to an operator of population dynamics, here we mean by the word ‘dimensionality reduction’. In life-cycle graph, the projection eliminates secondary groups and reallocate the reproductive outputs through secondary classes into primary classes (Figure 1).

Since the matrix 𝒫_𝕏_(*G*) (if well-defined) has a reduced dimension, it is appealing to utilize this matrix for obtaining invasion fitness. Indeed, in Appendix C, we show that 𝒫_𝕏_(*G*) exists whenever selection is weak; consequently,

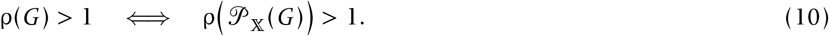

We refer to 𝒫_𝕏_(*G*) as projected next-generation matrix, or PNGM of *G*. The spectral radius of PNGM is referred to as the type-reproduction number in theoretical epidemiology (Roberts & Heesterbeek 2003).

The PNGM may be also derived using the original NGM framework. Given a decomposition of *J* = *U* − *V*, we consider another decomposition and the resulting NGM as follows:

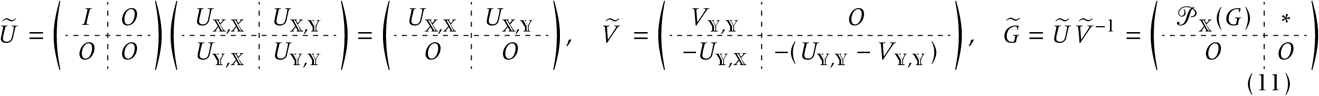

where the asterisk is some matrix, giving 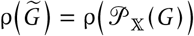. Hence, PNGM can be regarded as an outcome of NGM with different decomposition of the Jacobian.

### Properties of the framework

Beyond the invasion condition itself, the present framework possesses several, biologically useful properties (Box 3). First, the dimensionality reduction via the PNGM effectively separates the timescales among classes, where the eliminated (secondary) classes are assumed to equilibrate instantaneously relative to the retained (primary) classes (Appendix C). Second, the reproductive values of the retained classes are preserved (Appendix C). Finally, the framework preserves not only the invasion threshold, but also local stability properties around the evolutionary equilibrium, thereby enabling the analysis of long-term consequence of evolutionary dynamics (Appendix D and E).

### Summary of the framework

In practice, the invasion determinant and the PNGM serve complementary purposes within structural evolutionary invasion analysis. The invasion determinant yields a closed-form, scalar algebraic condition. This makes it optimal for analytical manipulation, differentiation, and evaluating convergence or evolutionary stability (Boxes 2 and 3). In contrast, the PNGM (Roberts & Heesterbeek 2003) structurally compresses the life-cycle graph while preserving reproductive-value weightings. It thus excels in biological interpretation, clarifying which pathways contribute most to mutant success. When applying the PNGM, we recommend designating key events (e.g., birth) as primary classes, leaving transient states as secondary. Because the true invasion threshold is invariant under these reductions, researchers can freely choose the partition that yields the most tractable and interpretable result.

## 4 Examples

We present four examples for illustration (Appendix E).

### Example 1: Two-class model

Consider a minimal non-trivial two-class model of two reproductive states (applicable to host-parasite or metapopulation dynamics). The mutant dynamics around the resident equilibrium read:

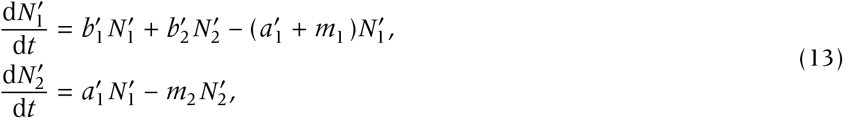

where *b*′s represent fecundity, which may be subject to density-dependent reduction; *a*′ represents transition; and *m* represents mortality (each with a subscript). The primed parameters *a* or *b*’s are evolving trait(s) though not necessarily independent.

Directly analyzing the Jacobian *J* underscores the difficulty of conventional approaches. For example, using Equation (6) yields:

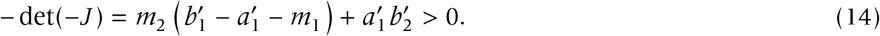

This inequality carries units of squared rates ([time^−2^]), obscuring its biological meaning.

To resolve this, we partition the Jacobian in Equation (13) as:

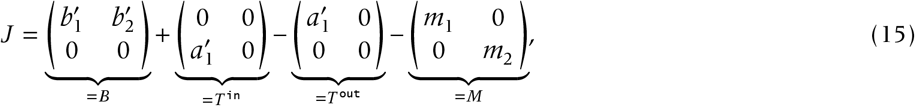

where from left to right, the matrices represent birth, transition-in, transition-out, and mortality. We write, dropping the prime for brevity, 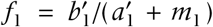 and 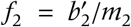 for the reproductive outputs in stage 1 and 2, and 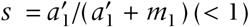 for survival probability.

Setting *U* = *B* + *T* ^in^ and *V* = *T* ^out^ + *M* yields the NGM:

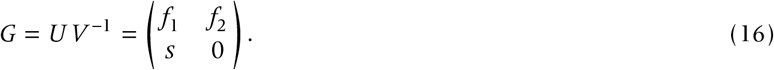

The spectral radius reads 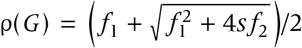, which is impossible to interpret (see Iritani & Cheptou 2017; Iritani *et al*. 2019).

The PNGM provides biological clarity. By choosing 𝕏 = {1}, we obtain the invasion condition:

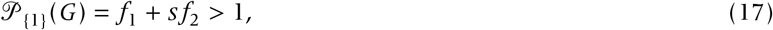

which rigorously recovers the biologically intuitive result that the lifetime reproductive success is a survival-weighted sum of total reproductive outputs.

While the invasion determinant yields the same invasion condition, the two-dimensional system has a special formula:

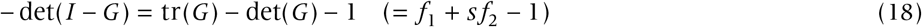

where tr() represents the trace (diagonals total). This simple determinant-trace formula holds only for the two-class model; there does not exist an analogous equation for higher dimensional systems.

While the PNGM is optimal for biological interpretation, the determinant form is particularly advantageous for algebraic manipulation. We derive the selection gradient (the first derivative of the invasion fitness with respect to the mutant phenotypic deviation δ). Using the technique in Box 3 and the neutrality condition *w*^◦^ = *f*_1_^◦^ + *s*^◦^ *f*_2_^◦^ = 1 yields:

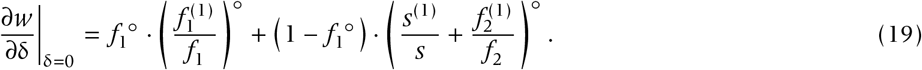

Equation (19) tells us that the life-time selection gradient is a weighted sum of the marginal changes in stage-specific fecundities. Specifically, the weights *f*_1_^◦^ and 1 − *f*_1_^◦^ represent the probabilities that an offspring is born to a class 1 and class 2 parent, respectively. An analogous analysis yields the second derivative required for assessing evolutionary stability, bypassing complex calculation and differentiation of the eigenvalues of the original NGM (Appendix D).

### Example 2: Evolutionary epidemiological model

The second example is the sex-structured evo-epidemiological model of Mitchell *et al*. (2022). While the authors have considered host-pathogen coevolution, we focus on host evolution for illustration.

The mutant dynamics reads:

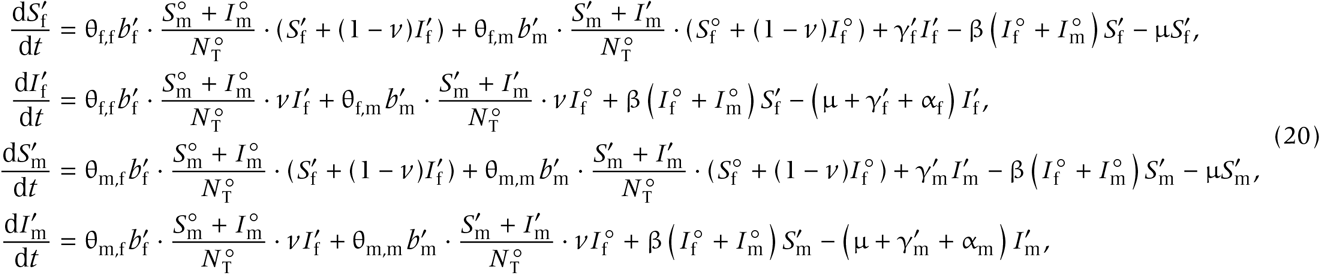

Where 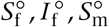, and 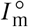 are the resident equilibrium densities of susceptible female, infected female, susceptible male, and infected male, respectively. In this ODEs, we write *b* for birth (subscript f for female and m for male), which is one of the evolving traits; θ for inheritance (e.g., all θs are 1/2 for diploids); *v* for vertical transmission of disease (only from female); γ for recovery, the other evolving trait; β for transmission rate; µ for natural death; and α for virulence.

To decompose the resulting four-by-four Jacobian, we put only birth terms (i.e., terms with coefficient *b*′s) into *U* while infection, recovery and mortality into *V* :

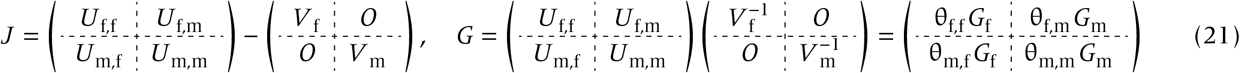

where *G*_f_ and *G*_m_ are fitness components of female and male, respectively, ignoring the genetic inheritance system.

We derive the invasion fitness for diploidy and haplodiploidy separately. NGM for diploidy reads:

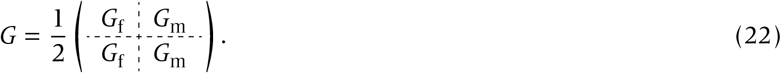

Crucially, because offspring receive equal genetic contributions from parents, NGM contains identical rows and has rank 2 (Mitchell *et al*. 2022, Eqn 10b). Thus, the invasion condition can be written as ρ(*G*_f_ /2 + *G*_m_ /2) > 1, i.e., the NGM is given by the arithmetic mean of sex-specific NGMs. Since *G*_f_ and *G*_m_ are both two-by-two, the two-class model results are readily applicable to obtain the explicit invasion condition.

In contrast, this arithmetic-mean formula breaks down for haplodiploidy (no fathers for males, θ_m,m_ = 0), making direct algebraic extraction of the spectral radius difficult. This is exactly where the PNGM becomes essential. Applying PNGM yields the manageable invasion condition ρ(*G*_f_ /2 + *G*_m_ *G*_f_ /2) > 1 (Appendix E). Biologically, this multiplicative term (*G*_m_ *G*_f_) captures the asymmetric nature of gene transmission: a male’s reproductive success is bottlenecked by his daughters’ fitness. Consequently, this structural derivation formulates a fundamental asymmetry among sexes (e.g., intralocus sexual conflicts). Our framework thus readily handles asymmetric genetic systems and its impacts on the evolution of sex-specific traits.

### Example 3: Discrete-time Lefkovitch model

We now consider discrete-time models and demonstrate how to perform the analogous analysis. The discrete-time mutant dynamics linearized around the resident equilibrium is given by:

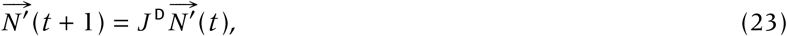

where *J* ^D^ (= *G*) represents the Jacobian (^D^ for ‘discrete-time’), and is nonnegative and irreducible. The invasion condition reads ρ(*J* ^D^) = ρ(*G*) > 1.

We analyze Lefkovitch’s (1965) model to illustrate the biological interpretation of PNGM. Consider one juvenile-stage (state 1) and three reproductive adult-stages (states 2, 3 and 4). We first construct the NGM and partition the classes as 𝕏 = {*X*_1_, *X*_2_, *X*_3_} and 𝕐 = {*X*_4_} (Figure 2):

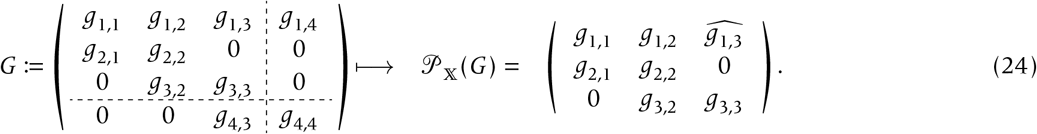

**Figure 2:**
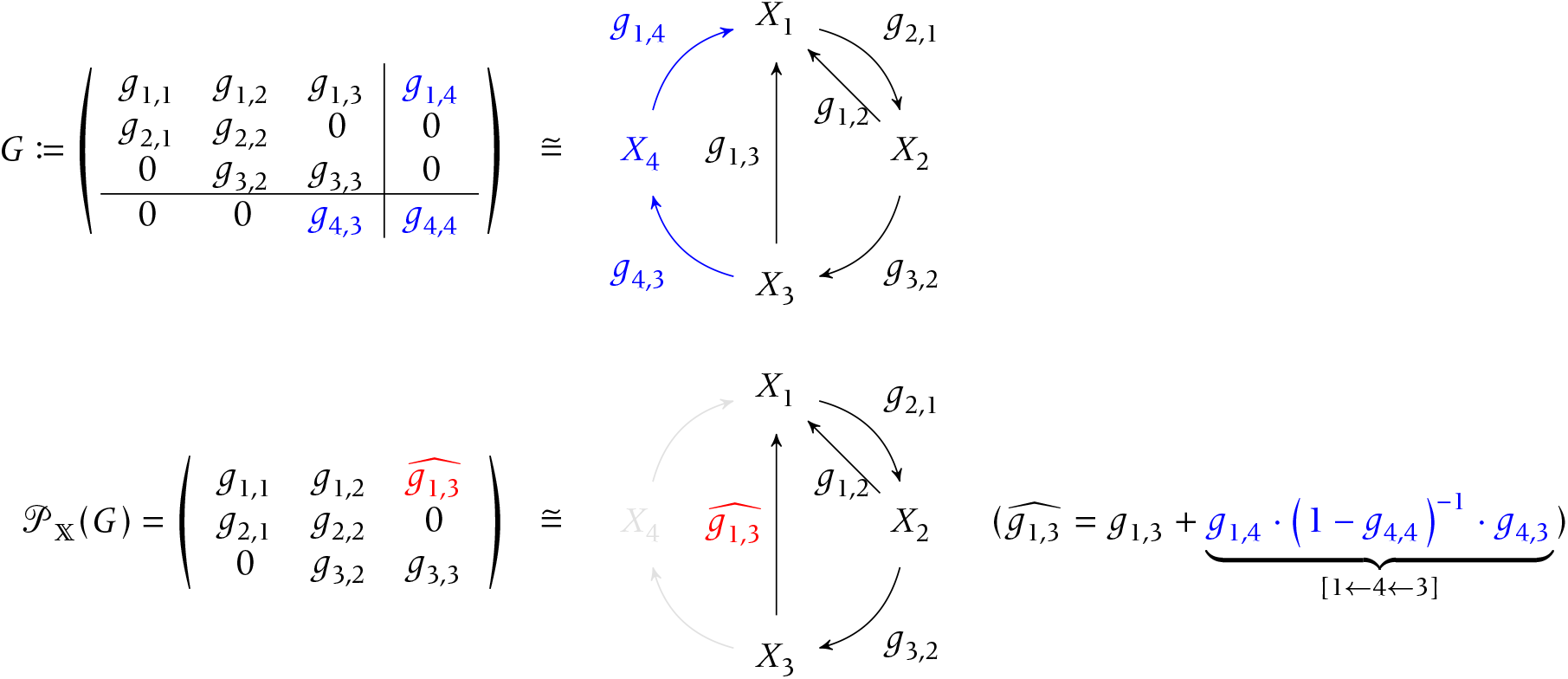
Lefkovitch model. Top: the original NGM *G* and corresponding life-cycle graph. Self-loops are not shown for simplicity. Bottom: PNGM and resulting life-cycle graph. We see *g*_1,3_ is changed to 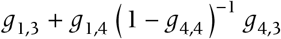, which means that the reproductive pathway from class-3 to class-1 now accounts for the reproductive pathway via class-4.

In this example, PNGM modifies the (1, 3)-rd element as 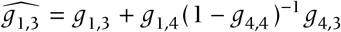, where the second additional term represents the reproductive success of an individual in secondary class 3 via class 4 (Figure 2).

The invasion condition (derived in Appendix E) reads:

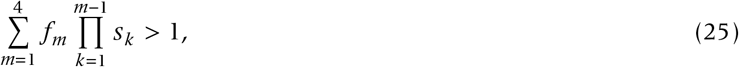

where *s*_*k*_ represents the survival probability of an individual in class *k*. Equation (25) represents the canonical form of lifetime reproductive success: the sum of stage-specific fecundities weighted by the cumulative probability of surviving to that stage. Rueffler & Metz (2013) (Theorem 3) detail the precise conditions under which reproductive success reduces to this canonical expression.

To further examine the long-term evolutionary outcomes, one may differentiate the left hand side of this invasion condition, using the techniques given in Box 2. The resulting formula yields a biologically transparent expression analogous to Fisher’s (1941) average excess, linking selection gradients to life-history trade-offs (e.g., survival versus reproduction; see also Giaimo & Traulsen 2019, 2021).

Crucially, our framework automatically resolves the algebraic hurdles of previous graph-theoretic methods (Hastings & Botsford 2006a; Rueffler *et al*. 2012; Lewis *et al*. 2019). For empirical ecologists leveraging large projection matrices from repositories like `COMPADRE` (Salguero-Gómez *et al*. 2014), the PNGM structurally compresses complex demographic data into tractable metrics. This enables researchers to apply empirical matrix demography to life-history evolution frameworks (e.g., Shefferson 2025).

### Example 4: Dispersal in stable habitats

We analyze the evolution of dispersal in a saturated, infinite-patch metapopulation, where each patch contains *K* reproductive adults (Hamilton & May 1977; Wright 1931). Demography follows a birth-death Moran process: an adult is replaced through random competition by a newborn offspring. Dispersal occurs at rate *d* with a mortality cost *c*.

Assume a mutant with dispersal rate *d*′ = *d* + δ arises under weak selection (|δ| ≪ 1). Let *k* ∈ {1, …, *K*} denote the number of mutants in a focal patch. The transition probability 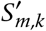 from *k* to *m* mutants follows (Ewens 2004):

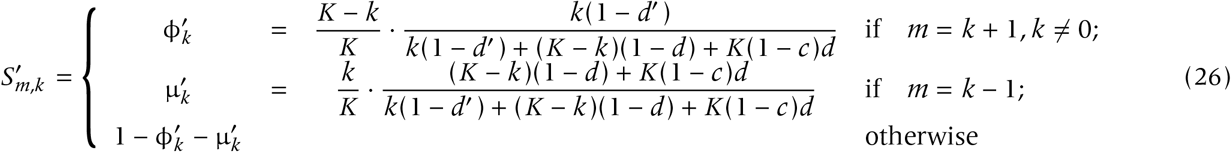

By convention, 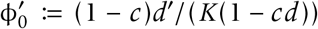 is the transition probability from zero to one mutant. Let 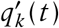 be the probability that a focal patch contains *k* mutants, conditional on containing at least one. The discrete-time dynamics for these patch states are:

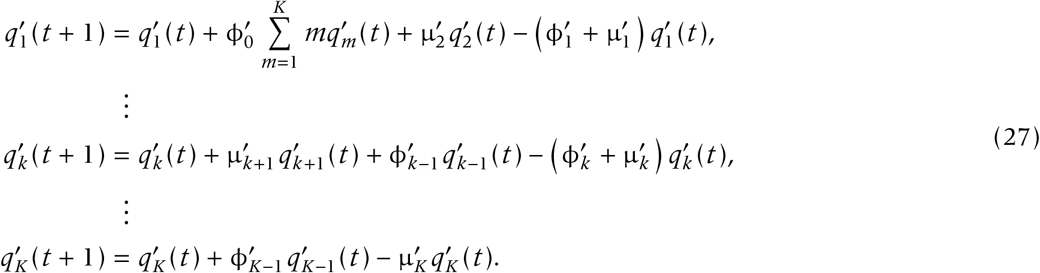

To derive the invasion condition, we recursively apply a quasi-equilibrium approximation to Equation (27). Setting 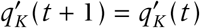 yields 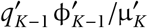. Substituting this backward and repeating the procedure recursively eliminates all intermediate states, yielding 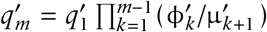 for *m* = 2, …, *K*. Substituting these into the first line of Equation (27) gives:

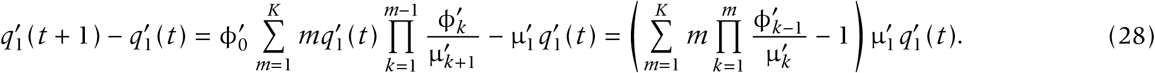

Thus, the invasion condition reduces to:

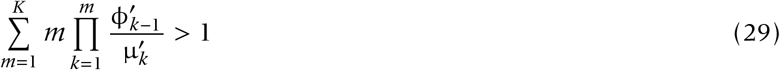

(Mullon & Lehmann 2014). This threshold reflects the metapopulation fitness concept (Metz & Gyllenberg 2001; Ajar 2003; Massol *et al*. 2009; Mullon *et al*. 2016) and closely resembles the Lefkovitch model (Equation (25)), with product terms representing expected sojourn times (Mullon & Lehmann 2014).

This derivation highlights two key points. First, it rigorously formulates inclusive fitness for spatially structured kin-selection models (Taylor & Frank 1996; Metz & Gyllenberg 2001). Second, recursive quasi-equilibrium approximations elegantly collapse high-dimensional, tri-diagonal systems (e.g., Moran processes) into tractable scalars (Ewens 2004; Sigmund 2010). While denser structures like the Wright-Fisher process complicate direct algebraic reduction (Ajar 2003), one can instead leverage matrix derivatives to evaluate stability conditions directly (Box 2; Appendix D).

## 5 Discussion

We have introduced “structural evolutionary invasion analysis”, a unifying framework for class-structured populations. By integrating the invasion determinant and the projected next-generation matrix (PNGM), this approach systematically simplifies complex evolutionary demography. The determinant yields tractable algebraic thresholds (Taylor & Bulmer 1980), while the PNGM clarifies biological pathways by formalizing timescale separation and preserving Fisher’s (1930) reproductive values (Box 3). As illustrated across our diverse examples, these combined tools provide a rigorous method for evaluating both short-term invasion and long-term evolutionary stability.

The invasion determinant provides a general algebraic tool to derive invasion conditions (Taylor & Bulmer 1980; Courteau & Lessard 2000; Rueffler *et al*. 2012). Our derivation relies crucially on the assumption of weak selection, where the phenotypic effect of a mutation is small. This assumption, however, is not highly restrictive; it often provides a robust approximation for evolutionary dynamics (Mullon & Lehmann 2014) and can be often relaxed (Metz & Leimar 2011). Furthermore, the invasion determinant facilitates the derivation of higher-order properties, such as the stability of evolutionary equilibria, and serves as the mathematical foundation for the negative decomposition theorem. Thus, while the bare determinant might sometimes lack immediate biological transparency, it constitutes the analytical core of structural evolutionary invasion analysis.

The projected next-generation matrix (PNGM) systematically simplifies life-cycle graphs by eliminating secondary classes. While this method builds upon the type-reproduction number theory from epidemiology (Roberts & Heesterbeek 2003), our synthesis adds two biological insights for evolutionary demography (Box 3). First, PNGM effectively separates the timescales of class dynamics. This enables researchers to place any subset of secondary classes into a quasiequilibrium state without requiring an actual timescale-difference (Example 4). (Morita *et al*. 2025) applied this method to infinite-dimensional systems and successfully analyzed the coevolution of floral sex allocation. By compressing these highly structured intermediate states, the PNGM transparently exposes the net reproductive pathways, as illustrated by the Lefkovitch model (Example 3). Second, the PNGM preserves Fisher’s (1930) reproductive values for the retained focal classes. In structured populations, variations in how different classes transmit alleles (nonheritable class transmission) complicate the evaluation of allele-frequency changes (Taylor 1990; Frank 1998; Rousset 2004; Priklopil & Lehmann 2020, 2024). Applying the structural reduction demonstrates that tracking gene-frequency changes within the focal subset is sufficient to determine mutant invadability.

Our structural approach naturally complements integral projection models (IPMs), which incorporate continuous traits (e.g., size, weight, or age) to capture ecologically realistic individual heterogeneity (Takada & Nakajima 1992; Caswell 2001; Rees & Ellner 2016; Giaimo 2025; Shefferson 2025). Although IPMs rely on infinite-dimensional integral operators rather than finite matrices, the core logic of the PNGM framework remains robustly applicable. Conceptually, the analogue of the PNGM in IPMs simply involves partitioning the continuous physiological space into discrete regions (Inaba & Nishiura 2008), practically collapsing the system back into a standard matrix problem. Furthermore, while our framework systematically reduces the dimensionality of class structures under the assumption of a monomorphic resident population, adaptive dynamics can ultimately lead to the stable coexistence of multiple lineages. To capture such polymorphic eco-evolutionary dynamics, our structural approach could be integrated with frameworks that approximate complex trait distributions, such as the oligomorphic theory (Sasaki & Dieckmann 2011; Lion *et al*. 2022, 2023; Wickman *et al*. 2023). Ultimately, this structural approach helps bridging the gap between rich life-history data and long-term eco-evolutionary dynamics.

Finally, we outline the underlying ecological assumptions for applying this framework. First, we assume that the ecological dynamics are regulated by density-dependence to reach a unique, strictly positive, and globally stable equilibrium (Cushing 1998; Geritz *et al*. 2002). A standard local stability analysis is typically sufficient to ensure that the resident equilibrium is stable. While bistability or multiple attractors can complicate the analysis, such systems can generally be handled by analyzing the invasibility of mutants at each stable equilibrium separately.

Second, we assume that the class-structure is irreducible (Stott *et al*. 2010). While seemingly restrictive, this assumption aligns with the premises of adaptive dynamics. In reducible models, gene flow between certain classes is permanently blocked, which contradicts the invasion-implies-substitution principle (Figure 1A). For instance, if a system has a one-way transition from class 2 to 3 with no return path (Figure 1A-i), a mutant arising in class 3 has no chance of fixation regardless of its selective advantage. Similarly, a population split into two disconnected subgroups (Figure 1A-ii) functions as two independent systems and must be analyzed as such. Thus, the irreducibility assumption simply ensures that the population forms a single, cohesive evolutionary unit.

To conclude, we have unified the methodology of evolutionary invasion analysis for class-structured populations. By integrating the invasion determinant and the projected next-generation matrix, we provide researchers with flexible tools that can be used interchangeably or in combination. Furthermore, this framework complements integral projection models, paving the way for the analysis of infinite-dimensional evolutionary demography. By systematically compressing complex demography into tractable metrics, this structural approach equips both theoreticians and empirical ecologists to bridge the gap between rich life-history data and long-term eco-evolutionary dynamics.

## 6 Acknowledgement

We thank Ryuichiro Isshiki and Takenori Takada for useful comments.

## Box 1 Classifying NGMs based on the invasion threshold

In the main text, we have seen that some NGMs have the same invasion condition. It is therefore useful to classify typical formulas of the NGMs with shared invasion threshold. In this box, we say that two NGMs *G*_1_ and *G*_2_ are invadability-equivalent if ρ(*G*_1_) > 1 ⇔ ρ(*G*_2_) > 1.

In Appendix C, we show that the following NGMs are invadability-equivalent, whenever *F* and *S* are both nonnegative regardless of their actual biological meanings: *F* + *S*, (*I* − *S*)^−1^ *F, F* (*I* − *S*)^−1^ and 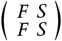 (Box-Fig 1). The last NGM has twice as large size as the original, and may be practically relevant when the class transition is unbiased (Rueffler *et al*. 2012; Lion & Metz 2018; Iritani *et al*. 2019, also see Example 2). We duplicated the focal class into two replicas with the same reproductive value. The life-cycle graph is expanded by creating a replica of the duplicated classes.

Whenever the inversion exists and *F* and *S* are nonnegative, these NGMs are invadability-equivalent to each other. This result is therefore a simple generalization of the so-called ‘fundamental matrix’ in population demography (Cushing 1998; Li & Schneider 2002).

One may wonder which NGM to use. One may preferably use the ‘snapshot’ fitness *F* + *S* for its numerical stability over the others involving matrix inversions. From an algebraic perspective, determining the stability properties of evolutionary equilibrium involves partial derivatives; again, the snapshot form (*F* + *S*) may seem preferable, but at the cost of higher dimensionality than the inverse formulas such as *F* (*I* − *S*)^−1^ if *F* contains many zeros or redundancies (said to be ‘rank deficient’ in linear algebra).

**Box-Fig S1:**
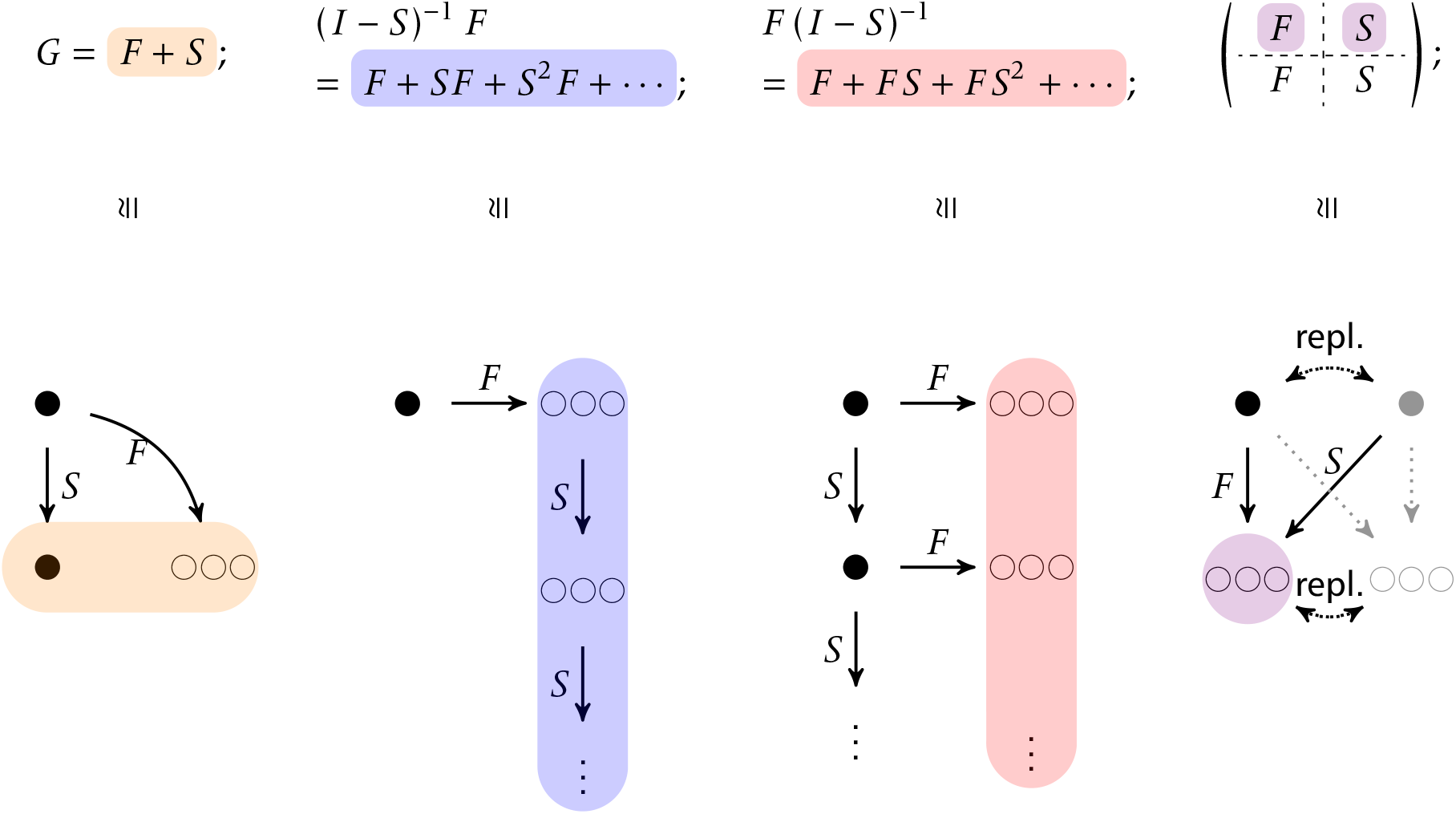
The invadability-equivalent NGMs (top) and the corresponding structures of NGMs (bottom). The colors correspond to the highlighted terms in the equation. Closed circle: focal individual; open cicles: offspring of focal individual; Gray circle: focal-individual replica. Gray pathway should not be counted in to avoid the double-counting of fitness. “repl.” for mirrored replica.

## Box 2 Evaluating selection gradients and life-history trade-offs

In practice, the invasion condition may involve fractional terms such as *w*′ := ϕ(*z* + δ, *z*)/µ(*z* + δ, *z*), where *z* is resident phenotype, with δ small. The derivative has a useful formula:

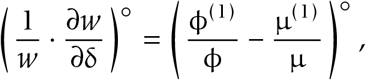

where ^(1)^ represents the first derivative with respect to δ. This is a form of the marginal value theorem linking the trade-off relationship between the marginal increases in the numerator (typically, fecundity) and the denominator (typically, mortality) (Charnov 1976). In population genetics, this expression is called as the average excess of gene substitution (Fisher 1941).

Invasion conditions often contain the cumulative survival probability terms like *s*′∏_*k*_*s*_*k*_ where *s*_*k*_ represents a survival or transition probability from class *k* and ∏_*k*_ represents the product operator over *k*. Its derivative also has a useful formula:

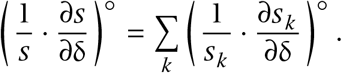

This expression can be also viewed as selection leading to the balanced, marginal increases in survival probabilities.

The common technique behind these derivatives is logarithmic differentiation to compute marginal changes in reproductive success. The following, matrix-analogue of the logarithmic derivative is also useful:

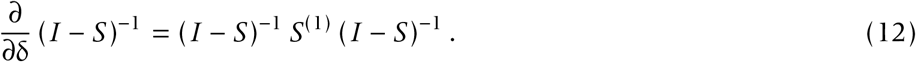

The matrix (*I* − *S*)^−1^, called as the fundamental matrix (Kemeny *et al*. 1966), represents the expected number of visits to a class starting from another class.

A note on interpretability: while symbolic computation software is powerful, automated simplification often obscures biological insight by expanding meaningful compound terms (e.g., life-history trade-offs). Differentiation should be viewed not merely as an algebraic operation but as a mapping that preserves the biological structure of fitness. Therefore, we recommend rearranging the derived equations to maintain biological interpretability, and prioritizing meaningful groupings over mathematical brevity.

## Box 3 Properties of the structural evolutionary invasion analysis

**Timescale separation** First, we obtain the PNGM as a result of separation of timescales between primary (which we assume is slow) and secondary (fast) groups. Biologically, this means that we can focus on only an arbitrary subset of classes to capture the overall gene-frequency changes (Appendix C). Note that one does not need any a priori knowledge or assumption of the actual differences in timescales of the classes. The utility of this method is clearly illustrated in the dispersal model (Example 3), where the class is defined as the number of mutant in a patch and the timescale is separated based on the purpose of deriving fitness, regardless of biological assumption on the timescales.

**Fisher’s reproductive values** Second, Fisher’s (1930) reproductive values of the primary group do not change with PNGM (Appendix C). This property reflects the reproductive values’ nature of measuring the asymptotically long-term contributions to the gene pool (Taylor 1990; Caswell 2001; Rousset 2004). More precise detail is encapsulated in Appendix C.

**Inheriting stability characteristics** Practically, the ultimate goal of applying the adaptive dynamics framework is to locate and characterize evolutionary equilibrium (EE), namely the direction of selection and possibility of disruptive selection. These characteristics of EE can be analyzed through:

i. the first derivative of invasion fitness (i.e., selection gradient capturing the location of EE);
ii. how the direction of selection changes with the resident trait value (i.e., convergence stability or attainability); and
iii. the second derivative of invasion fitness (i.e., whether selection is disruptive or stabilizing around EE; Maynard Smith & Price 1973; Maynard Smith 1982; Christiansen 1991; Takada & Kigami 1991; Eshel 1996).

Regardless of using the original NGM, invasion determinant, or PNGM, the resulting invasion conditions give the same stability properties (i)–(iii). Thus, one can characterize both the location and stability of EE using the structural evolutionary invasion analysis. See Appendix D for details.

## Appendix

### A Invasion determinant formula

#### Resident NGM

We here show that the resident NGM has a unity spectral radius. This result is obvious for discrete-time models. We thus derive the result for continuous-time models.

To see this, consider two types each with density 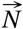 and 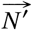. Ignoring mutation, we can describe their dynamics as:

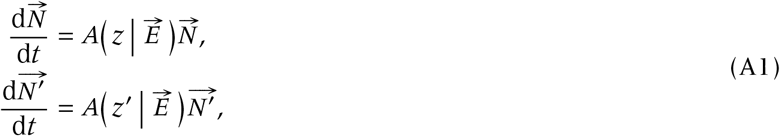

where 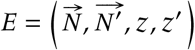. Note that these are generally nonlinear because of 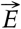.

At the resident equilibrium with no mutant,

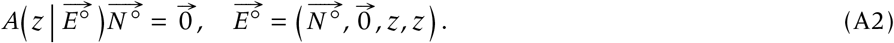

When the mutant is rare, its dynamics can be linearized as:

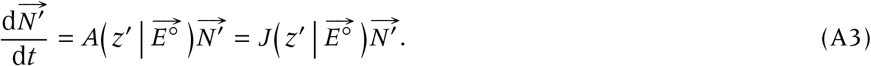

Thus, the Jacobian matrix evaluated at the resident equilibrium is exactly 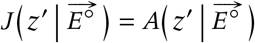.

At neutrality (*z*′ = *z*), substituting this into the resident equilibrium condition yields:

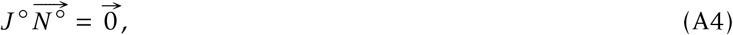

where 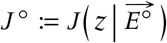.

Decomposing the Jacobian as *J* ^◦^ = *U* ^◦^ − *V* ^◦^ according to the NGM framework, we get:

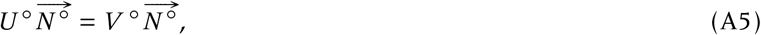

to which we pre-multiply *G*^◦^ = *U* ^◦^ (*V* ^◦^)^−1^ to get:

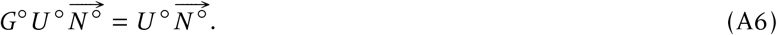

Because the matrix *U* ^◦^ is non-negative and the equilibrium density 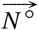 is strictly positive, the vector 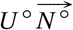 must be non-negative and non-zero. The equation 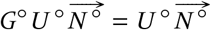 indicates that 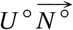 is an eigenvector of *G*^◦^ associated with the unit eigenvalue. By the Perron-Frobenius theorem for non-negative irreducible matrices, an eigenvalue associated with a non-negative eigenvector must be the spectral radius. Therefore, ρ(*G*^◦^) = 1.

#### Theorem on the invasion determinant

Throughout the appendix, we write “a ⋚ c ⇔ b ⋛ c” to mean that *a* is larger, equal, or smaller to *c* if and only if *b* is. We here prove the invasion determinant.

##### Theorem 1

(Invasion determinant formula). *Let G be a next-generation matrix, and assume that the selection is weak. Then*,

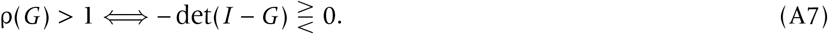

*Proof*. Let Γ_*M*_ (*x*) := det(*x I* − *M*) be the monic characteristic polynomial of a matrix *M*, and *G*^◦^ be the mutant NGM *G* evaluated at δ = 0 (i.e., the neutral mutant matrix). Thus, we have ρ(*G*^◦^) = 1 and so, by continuity of the spectrum of *G*, we have ρ(*G*) = ρ(*G*^◦^)+ε = 1+ε where ε is small. Furthermore, the Perron-Frobenius theorem for an irreducible nonnegative matrix implies that the spectral radius of *G* is a simple eigenvalue of *G* and so Γ_*G*_ (ρ(*G*)) = 0 and dΓ_*G*_ (*x*)/d*x*|_*x*=ρ_(_*G*_) > 0. As a result, if δ is small we can expand the equation Γ_*G*_ (1 + ε) = 0 in powers of ε to obtain Γ_*G*_ (1) + dΓ_*G*_ (*x*)/d*x*|_*x*=ρ(*G*)_ ε+ *o*(ε) = 0. Given ε = ρ(*G*) − 1, for small enough δ and thus ε this can be re-written as sign(ρ(*G*) − 1) = sign(−Γ_*G*_ (1)). Finally, from the definition of Γ_*G*_ (*x*) this can be re-written as sign(ρ(*G*) − 1) = sign(− det(*I* − *G*)) from which the theorem follows; Figure 3 provides an alternative, graphical proof.

**Figure 3:**
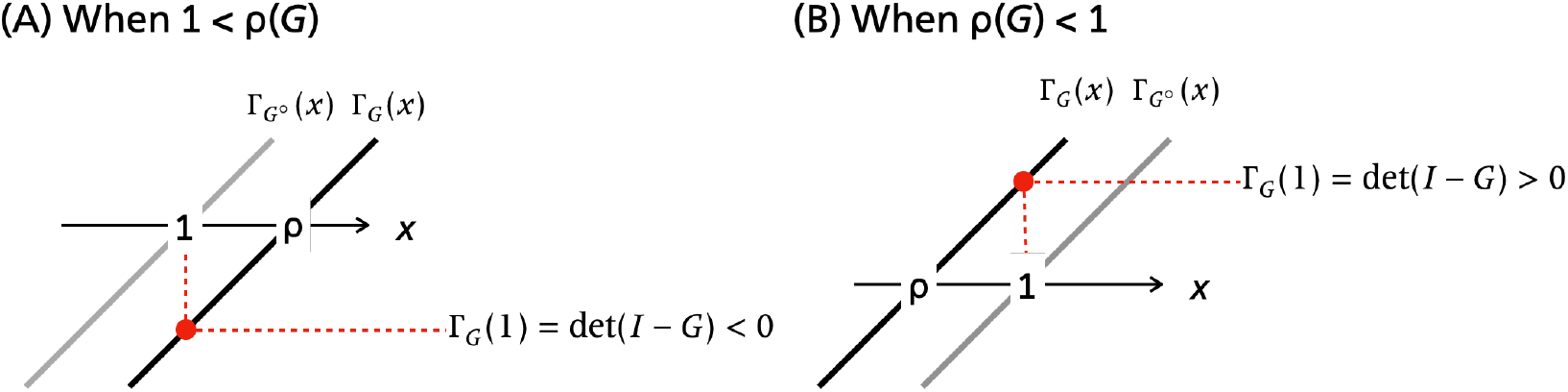
A graphical proof of Theorem 1.

The corresponding result for the original continuous-time Jacobian reads:

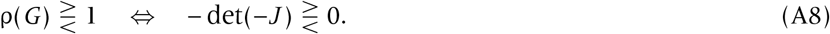

To prove this we decompose *J* = *U* −*V* into the components that satisfy the hypotheses of NGM. With some rearrangement, we get:

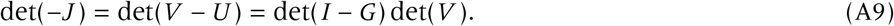

We notice that *V* has nonnpositive elements at off-diagonal components while positive element at diagonal components, and *V* ^−1^ is nonnegative. Using the theory of M-matrix, we have det(*V*) > 0 (Berman & Plemmons 1987). Therefore, det(−*J*) and det(*I* − *G*) have the same sign, and thus both can be used for invasion condition.

### B Nonnegative decomposition theorem

We here show that any NGM can be split into nonnegative and nonzero matrices and converted to an alternative, invadability-equivalent NGM.

#### Theorem 2

(Nonnegative decomposition theorem). *Let G be a (irreducible) next-generation matrix, and assume that the selection is weak. Put G* = *H*_1_ + *H*_2_ *where H*_1_ *and H*_2_ *are both nonnegative and nonzero for values of* δ *within some neighbourhood of* δ = 0. *Then*,

1. *for small enough* δ, ρ(*H*_2_) < 1 *so that the matrix I* − *H*_2_ *is invertible and thus* 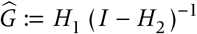 *is well-defined, and*
2. *if* 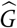 *is irreducible, the following equivalency holds:*

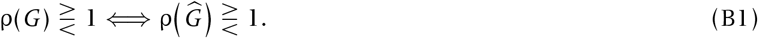

*Proof*. We first show that *I* − *H*_2_ is invertible whenever *H*_1_ is nonzero. For this, we use Corollary 2.1.5 of Berman & Plemmons (1987, p. 27), which is stated as follows:

#### Lemma 1

(Berman and Plemmons 1994). *Let A and B be nonnegative matrices. If B* − *A is nonnegative with B* − *A* /= *O, and if A* + *B is irreducible, then* ρ(*A*) < ρ(*B*).

We take 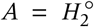 and *B* = *G*^◦^. We have 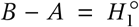, which is nonnegative and nonzero by our assumption. Furthermore, because *B* = *G*^◦^ is irreducible by the standard assumption of the NGM, and *O* ≤ *A* ≤ *B*, the sum *A* + *B* shares the exact same zero-nonzero pattern as *B*, making *A* + *B* irreducible as well. Thus, the hypotheses of Lemma 1 are satisfied, yielding 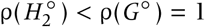. By the continuity of the spectral radius with respect to matrix entries, ρ(*H*_2_) is continuous in δ. Therefore, ρ(*H*_2_) < 1 under weak selection (i.e., for small enough |δ|), which implies that *I* − *H*_2_ is invertible and completes the proof of the first statement.

To prove the second part, we use the invasion determinant. By definition of 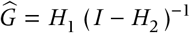, we have:

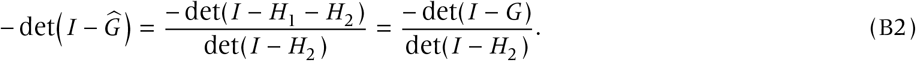

Because ρ(*H*_2_) < 1 and *H*_2_ is nonnegative, the monic characteristic polynomial of *H*_2_ is positive at unity, so 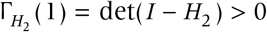. Therefore, 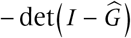 and − det(*I* − *G*) share the exact same sign.

Because *G* is irreducible, Theorem 1 guarantees that ρ(*G*) ⋚ 1 ⇔ − det(*I* − *G*) ⋛ 0.. Likewise, because the reduced matrix 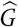 is explicitly assumed to be irreducible in the theorem statement, applying Theorem 1 to 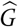 yields 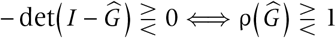. Connecting these equivalences through the shared sign of their determinants completes the proof.

The nonnegative decomposition theorem is based on a matrix splitting method known as ‘regular splitting’ for numerical analysis (Varga 1963).

### C PNGM

#### Derivation of PNGM

We here prove the following theorem.

##### Theorem 3

(Projected Next-Generation Theorem). *Let G be a next-generation matrix of a mutant, with the following block-partition structure:*

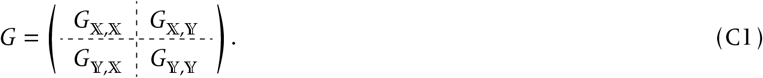

*Then, under weak selection, the following results hold true:*

1. ρ(*G*_𝕐,𝕐_) < 1.

*2*.

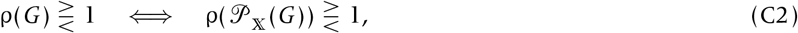

*where*

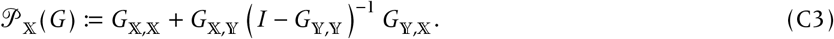

The first part can be proved in the same way as the first part of Theorem 1. The second part is also a direct implication of Theorem 2. In Theorem 2, we choose the decomposition as *H*_1_ := Π_𝕏_ *G* with *H*_2_ = *G* − *H*_1_ = Π_𝕐_ *G*, where Π_𝕏_ and Π_𝕐_ are projection matrices (Roberts & Heesterbeek 2003):

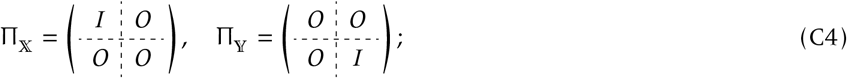

note that a matrix Π is called as projection if Π^2^ = Π (in mathematical sense).

We here provide a similar, alternative proof. Specifically, we use the Schur complement and Schur’s determinant identity (Crabtree & Haynsworth 1969).

##### Definition 1

(Schur complement). Let *M* be a square matrix block-partitioned as:

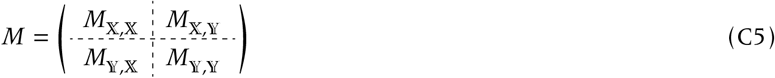

Suppose *M*_𝕐,𝕐_ is invertible. We define the Schur complement of *M* to *M*_𝕐,𝕐_, denoted by 𝒮_𝕏_(*M*), as

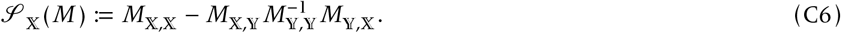

By definition, 𝒫_𝕏_(*M*) = *I* − 𝒮_𝕏_(*I* − *M*).

##### Lemma 2

(Schur). *Let M be a square matrix satisfying the assumptions of Definition 1*. *Then, the following identity holds true:*

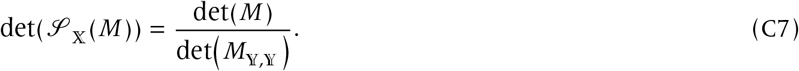

*(Proof of Lemma 2)*. Consider the identity:

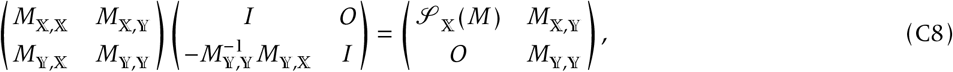

taking the determinant of which yields:

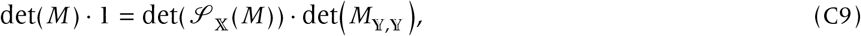

which yields the desired result.

*(Proof of Theorem 3)*. In Lemma 2, put *M* = *I* − *G*. By definition, we have:

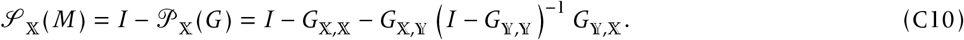

Applying Lemma 2, we have:

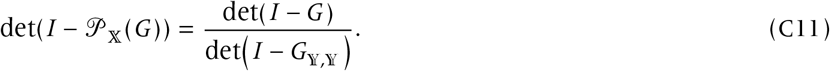

Because ρ(*G*_𝕐,𝕐_) < 1, the characteristic polynomial of *G*_𝕐,𝕐_ must be positive at unity, implying det(*I* − *G*_𝕐,𝕐_) > 0. Hence the LHS and the numerator of the RHS have the same sign. This completes the proof.

##### Interpretation of PNGM

We consider a mutant individual residing in one of the primary group classes and assess how many offspring it produces to the primary group. Specifically, we write *P*_*n*_ for a matrix of the total expected number of the mutant’s offspring produced to the primary group over *n* subsequent generations. We then get:

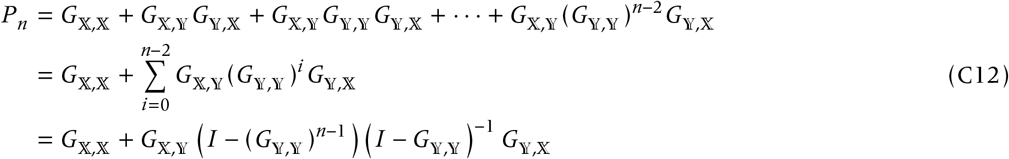

where the first term represents the number of primary group-offspring directly produced by the mutant in one generation; the second term represents the number of primary group-grand-offspring whose parents are in the secondary group (the remaining terms can be read analogously). Because ρ(*G*𝕐,𝕐) < 1, we have lim_*n*→∞_ *P*_*n*_ = 𝒫_𝕏_(*G*) (Horn & Johnson 1985, Theorem 5.6.12). Hence, the PNGM counts the total number of offspring in the primary classes produced by the mutant from the primary classes throughout the lifespan.

#### Embedding of PNGM

Here we examine the relation between PNGM and nonnegative decomposition theorem. We specifically consider embedding *G* = *H*_1_ + *H*_2_ into a high dimensional space as:

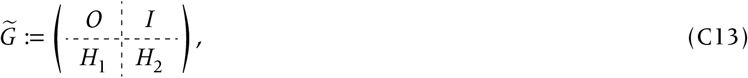

with:

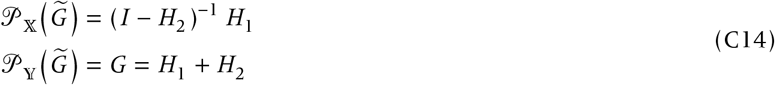

(Figure 4). On the other hand, we also have

**Figure 4:**
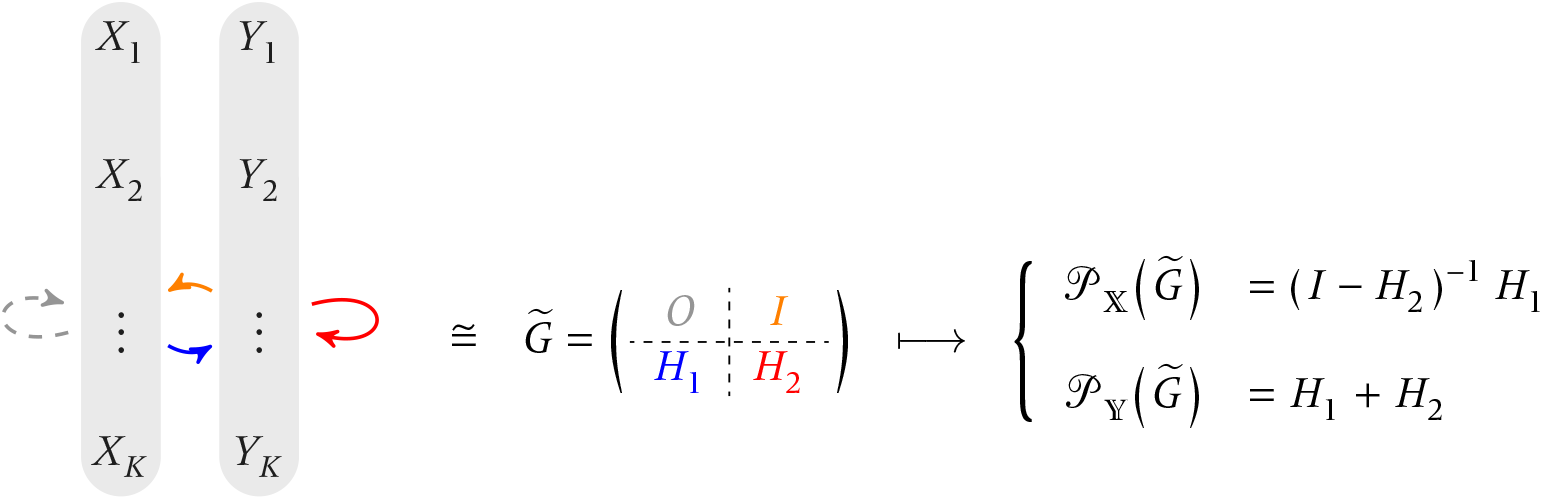
Embedding of *G* and its projection. Here, *Y* s are the orignal classes and *X* are their ‘replicas’.

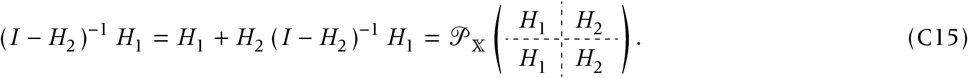

We note that *H*_1_ (*I* − *H*_2_)^−1^ and (*I* − *H*_2_)^−1^ *H*_1_ have the same spectral radius; indeed, because (*I* − *H*_2_)^−1^ is an invertible matrix, these two matrices are related by a similarity transformation:

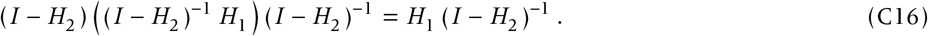

Because similar matrices share the exact same characteristic polynomial (and thus the exact same eigenvalues), their spectral radii are strictly identical (e.g., Horn & Johnson 1985). Thus, the following matrices share the same invasion conditions:

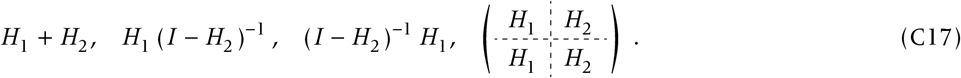

#### The relation to separation of timescales

We here show that the separation of timescales among classes produces the PNGM.

##### Theorem 4

(Constructing PNGM from continuous-time mutant dynamics). *Consider the following, K-dimensional mutant dynamics:*

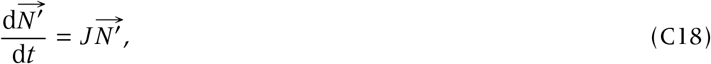

*where* 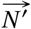 *represents a vector collecting the densities in the classes, and J represents the Jacobian, block-partitioned as Equation* (C1). *Consider the Schur-complement* 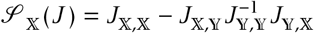. *Then*,

1. *taking the quasi-equilibrium approximation* 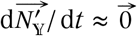 *leads to* 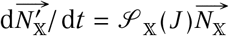;
2. *there exists a decomposition of J* = *U* − *V that satisfies the hypotheses of NGM theorem, V*_𝕏,𝕏_ *nonsingular, and* 𝒮_𝕏_(*J*) = 𝒫_𝕏_ (*U V* ^−1^)*V*_𝕏,𝕏_ − *V*_𝕏,𝕏_. *Thus for such U, V with U V* ^−1^ = *G, the resulting NGM is* 𝒫_𝕏_ (*G*)*V*_𝕏,𝕏_ (*V*_𝕏,𝕏_)^−1^ = 𝒫_𝕏_ (*G*).

*Proof*. (1) Let us calculate the Schur-complement for the original *K* -dimensional Jacobian. Consider the quasiequilibrium approximation given by:

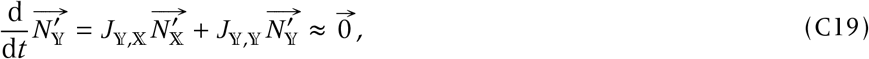

which gives 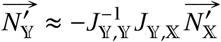, and thus

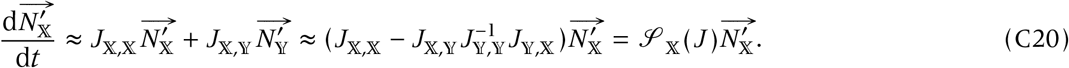

(2) It is easy to check that if we choose a block-diagonal, invertible, and inverse-nonnegative matrix as *V*, the theorem immediately follows.

This theorem guarantees the equivalence of quasi-equilibrium approximation (sepataion of timescales) and PNGM: the primary group dynamics are taken as ‘slow’ while the secondary group dynamics are ‘fast.’

We can consequently derive the discrete-time analogue of the separation of timescales.

**Proposition 1** (Constructing PNGM from discrete-time mutant dynamics). *Suppose the mutant dynamics:*

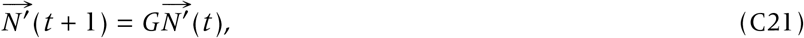

*where G represents a next-generation matrix of the system, block-partitioned as Equation* (C1). *Then, the PNGM obtains by taking the quasi-equilibrium approximation of slow-primary group and fast-secondary group*.

*Proof*. We explicitly solve 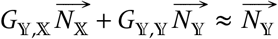 to get 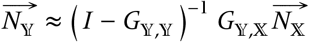. Substituting this into the dynamics of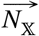 yields

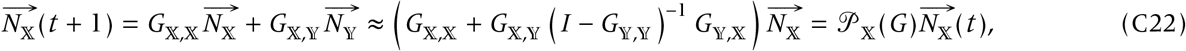

which proves the proposition.

#### Reproductive values

Here we show that the eigenvectors for the primary group at neutrality are invariant with the operation of projection, 𝒫_𝕏_. Let 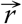 denote the right eigenvector of *G*^◦^ (at neutrality) associated with ρ(*G*^◦^) = 1:

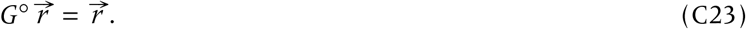

Now suppose that the vector is partitioned so the dimension is consistent with the block-partitioning of *G*^◦^; namely:

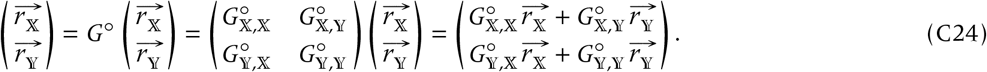

By solving for 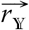 the linear equation at the second block, we have:

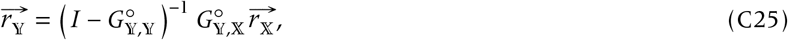

substituting which into the first block gives:

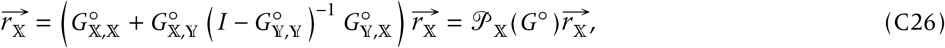

as desired. The same argument applies to the left eigenvector associated with the spectral radius ρ(*G*^◦^) = 1.

### D Stability analysis

We have presented various invadability-equivalent NGMs with κ _1_ (δ, *z*) := ρ(*G*_1_) − 1 ⋛ 0 if and only if κ _2_ (δ, *z*) := ρ(*G*_2_) − 1 ⋛ 0. From this expression, we can derive three properties of long-term evolutionary dynamics, namely (i) the location of evolutionary equilibrium (EE), (ii) convergence stability, and (iii) evolutionary stability.

We write sign() for the sign function.

#### Location of evolutionary equilibrium and convergence stability

We expand κ_*i*_ (δ, *z*) to the first order in δ as:

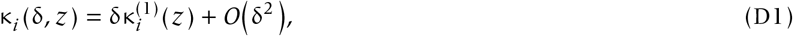

where 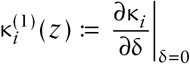 is the selection gradient, and the zero-th order term vanishes due to neutrality (κ_*i*_ (0, *z*) = 0).

For any given resident trait *z*, κ_1_ (δ, *z*) and κ_2_ (δ, *z*) share the same sign for sufficiently small |δ| > 0. For this point-wise property to hold, their leading-order coefficients in the Taylor expansion evaluated at *z* must also share the same sign. Thus, we have:

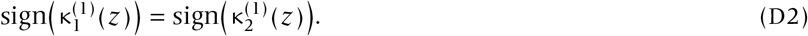

From this equivalence, two properties immediately follow: 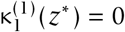 if and only if 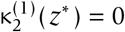, which proves that the location of the evolutionary equilibrium (EE) is identical. (ii) In the neighborhood of the EE, the sign of the selection gradient is identical for both measures. This implies that the direction of trait substitution is exactly the same, thereby ensuring that their convergence stability agrees.

#### Evolutionary stability

Fix the resident trait at an evolutionary equilibrium, *z* = *z*^*^. We expand the fitness functions with respect to δ up to the second order:

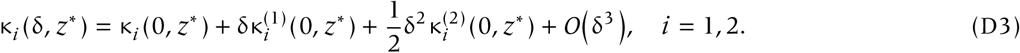

The zero-th order term vanishes due to neutrality (κ_*i*_ (0, *z*^*^) = 0), and the first-order term vanishes because *z*^*^ is an evolutionary equilibrium 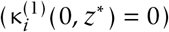.

Because κ _1_ (δ, *z*\*) and κ_2_ (δ, *z*\*) share the same sign for any sufficiently small |δ| > 0, their leading non-zero terms in the expansion must also share the same sign. Thus, we conclude that

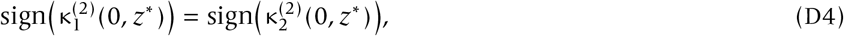

which ensures that the condition for evolutionary stability is identical between any two invadability-equivalent NGMs.

#### Reproductive value-weighted selection gradient

We show that the selection gradients derived either by the original NGM and PNGM agree. Let us derive the selection gradient using the reproductive values: we write 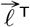 or 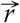 for the left or right eigenvector of neutral NGM *G*^◦^ associated with the spectral radius 1, respectively (Taylor & Frank 1996; Frank 1998; Avila & Mullon 2023). Given PNGM, we partition the eigenvectors according to the same partition-size:

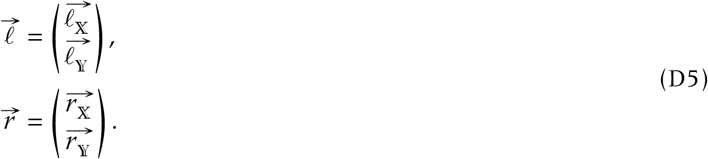

The first derivative of the PNGM, with the following lines being evaluated at neutrality *z*′ = *z* unless otherwise stated, is given by:

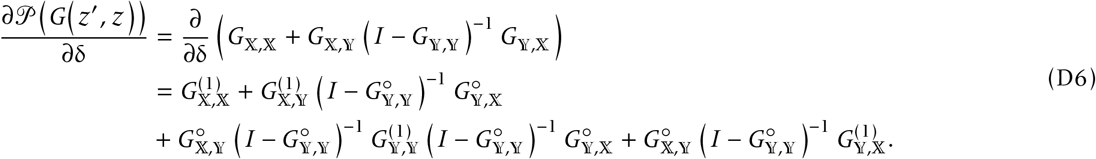

By definition, we notice:

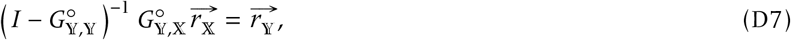

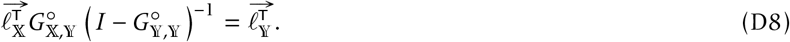

Premultiplying the vector 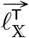 and also postmultiplying 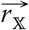 to the derivative gives:

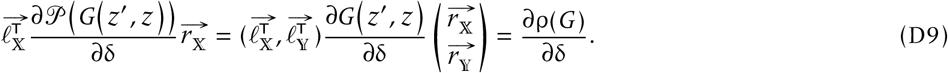

We have therefore confirmed the equivalence between the convergence stability conditions for the original and projected next-generation matrix.

### E Analyses of the examples

#### Example 1: Two-class model

We describe the result of NGM by taking the decomposition of *J* as in the main text:

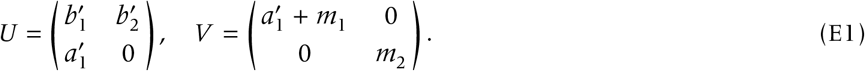

We write 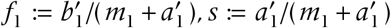 and 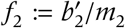 for the fecundity of class-1 individual, survival probability (0 ≤ *s* ≤ 1), and fecundity of class-2 individual, respectively. A direct calculation then gives:

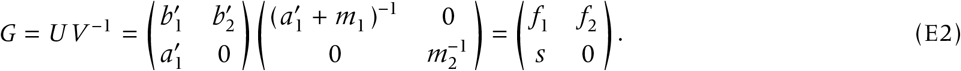

Using the invasion determinant yields the invasion condition, of:

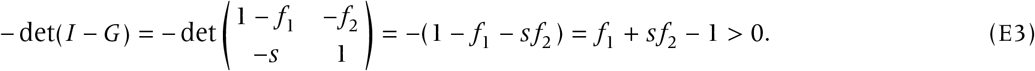

We thus get the invasion condition *f*_1_ + *s f*_2_ > 1 as derived in the main text.

#### Example 2: Epidemiological model

The mutant dynamics is given by:

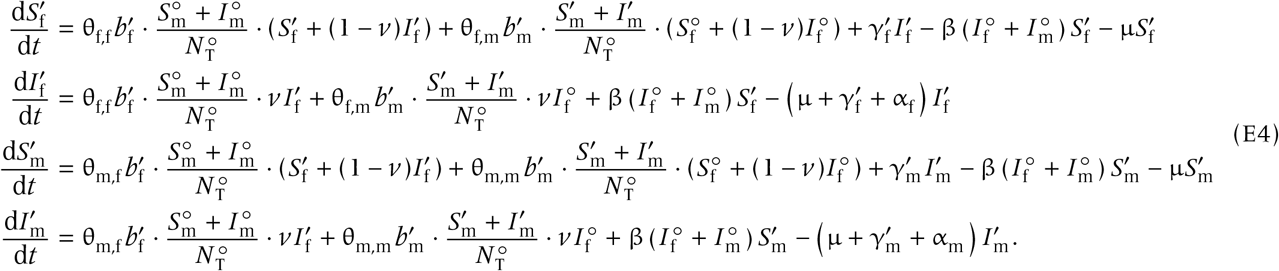

The resulting Jacobian reads:

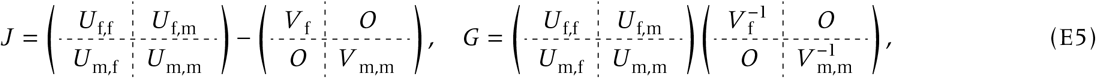

where, writing 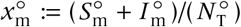 for the sex ratio, 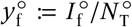 for the proportion of infected females, and θ_*k,m*_ for the probability that an individual of sex *k* derives a gene from a parent of sex *m* (for diploid organisms, these are all 1/2):

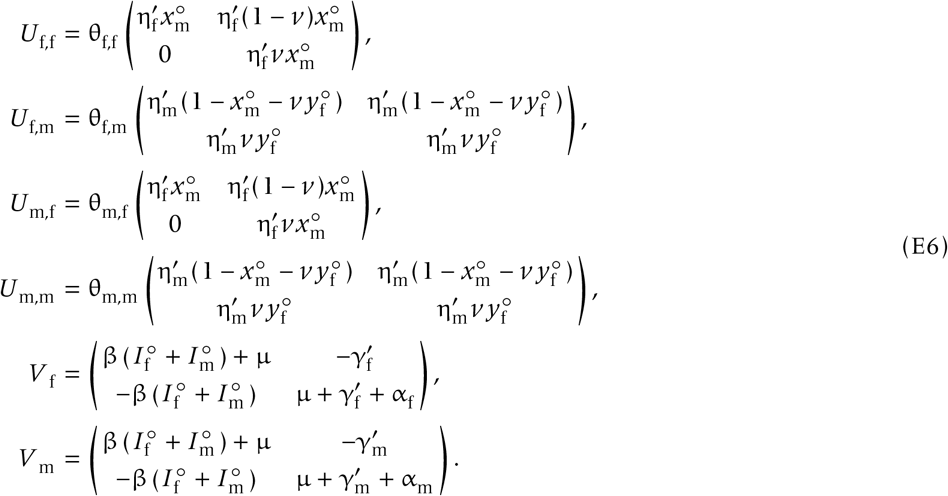

Rather than directly analyzing the four-by-four Jacobian, we use tensor decomposition to exploit biological symmetries. This elegantly compacts the system into a two-by-two form, facilitating numerical implementation:

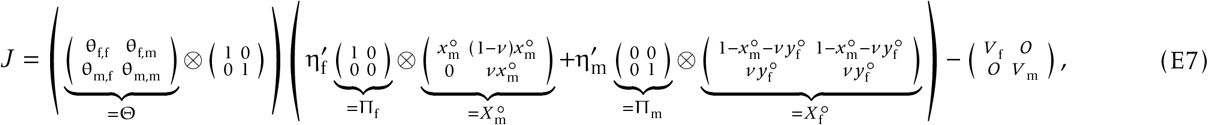

which, using the mixed-product rule (Graham 2018), we can rewrite as:

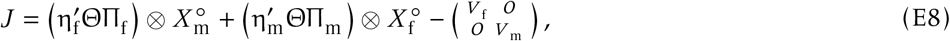

where Πs are projections:

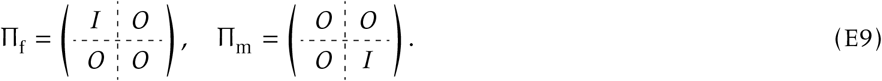

We invert the last *V* -matrix and get NGM:

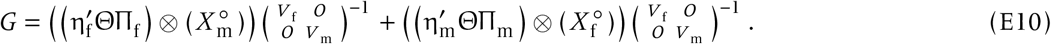

Using Π_f_ and Π_m_, we can rewrite the inversion as:

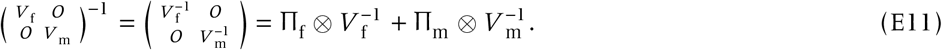

Again using the mixed product rule and 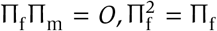 and 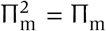, we get:

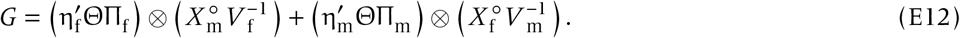

We note that, by writing 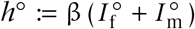,

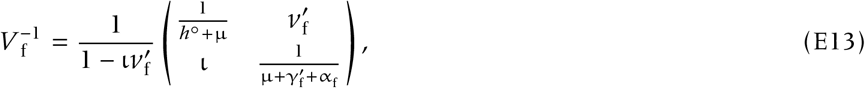

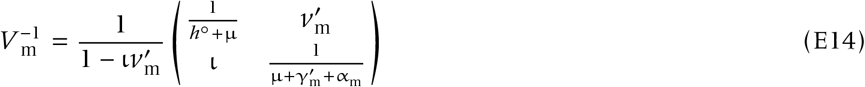

with 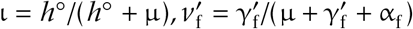 and 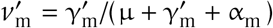. Taken together,

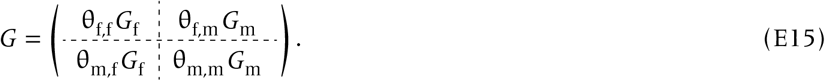

For diploidy,

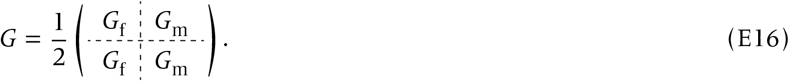

Since this takes the form of block-partitioned matrix in Box 1, the invasion condition reduces to:

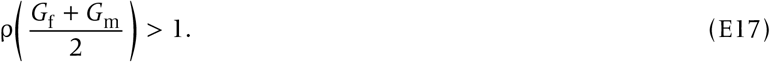

For haplodiploidy,

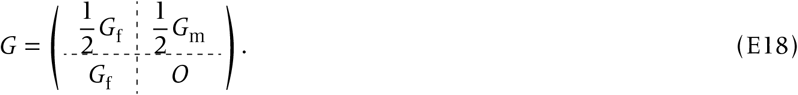

By applying the PNGM to eliminate the male class, we get:

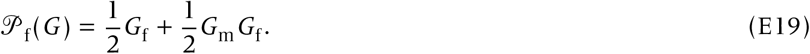

The first component represents the direct transmission of genes from female to female, and the second component represents the indirect transmission from female to female via male. The invasion condition for haplodiploidy is thus

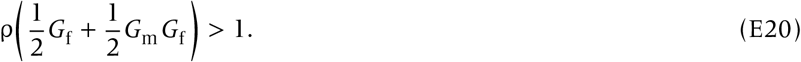

#### Example 3: Lefkovitch model

We consider the following NGM:

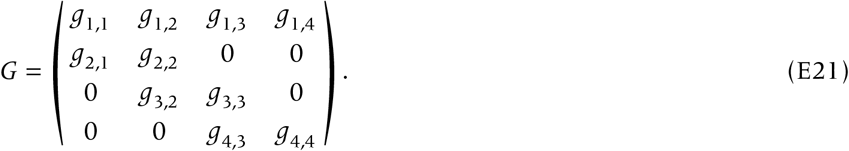

This NGM does not have a closed form of spectral radius. We partition the classes into primary and secondary groups: 𝕏 = {1, 2, 3} and 𝕐 = {4}. The PNGM then reads:

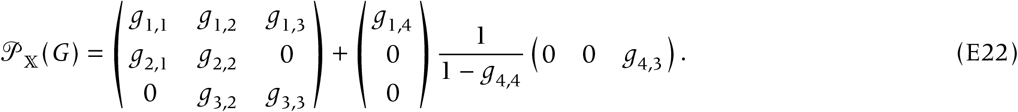

The second matrix is zero except for the (1, 3)-rd element *g*_1,4_ (1 − *g*_4,4_)^−1^ *g*_4,3_, which recovers the result of the main text.

To derive the ‘canonical form’ invasion condition ∑_*k*_ *f*_*k*_ ∏_*m*_ *s*_*m*_ > 1, we first partition the NGM into fecundity and transition components *G* = *F* + *S*, with:

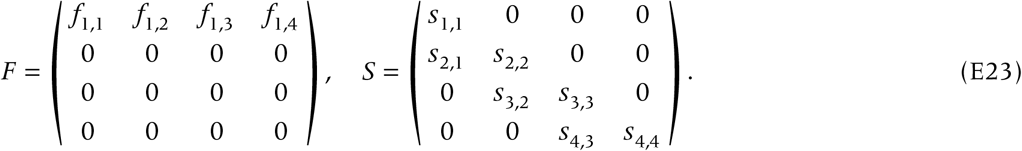

Inverting *I* − *S* yields:

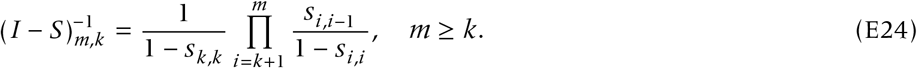

Since *F* has elements only in the first row, what we actually need is

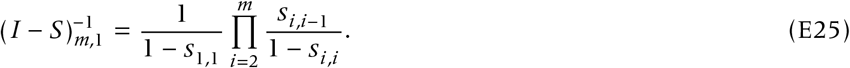

Using this we get:

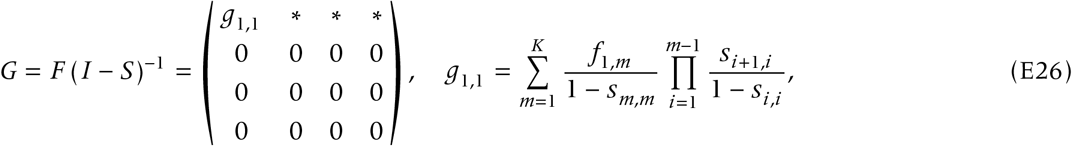

which thus says that ρ(*G*) = *g*_1,1_, leading to the canonical form given in the main text following the transformation:

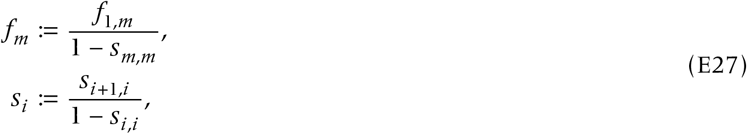

by abuse of notation.

We can derive the same canonical form by recursively applying 𝒫_{1,2,3}_, 𝒫_{1,2}_, 𝒫_{1}_ :

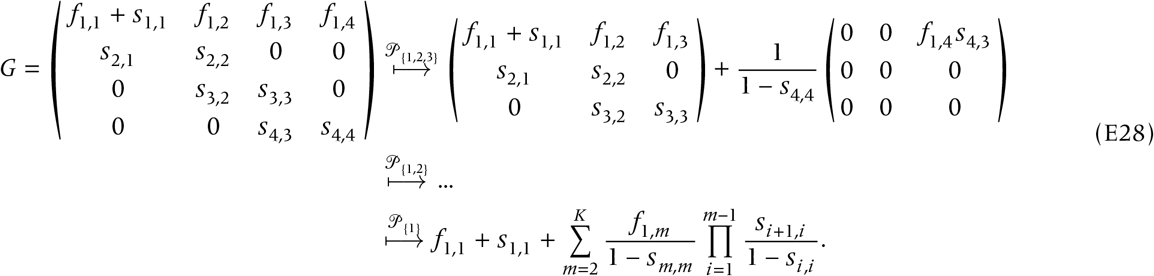

Rearranging the last quantity > 1 yields the canonical form of invasion condition.

#### Example 4: Dispersal in stable habitats

##### Moran process

The transition probabilities *S*_*k,m*_ are given by Equation (26) (e.g., Mullon & Lehmann 2014). Based on these probabilities, we can describe the dynamics of the number of mutants per patch as:

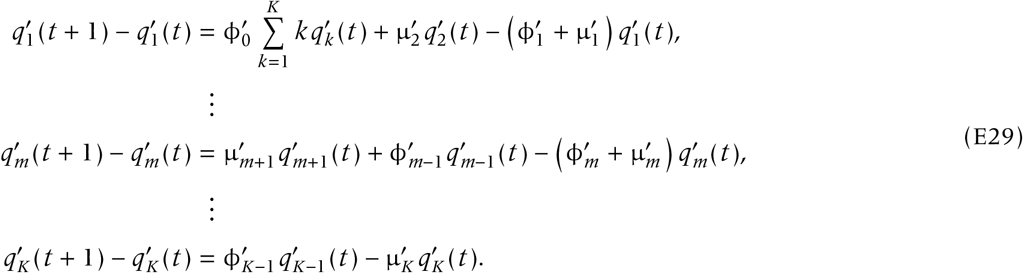

We now perform the separation of timescales. We eliminate the state variables in descending order from 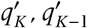, to 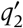. First, setting the left-hand side of the last line to zero yields 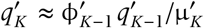. Second, we consider

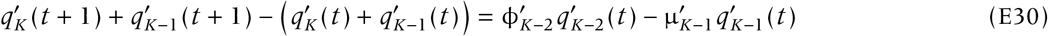

and set it to zero to obtain

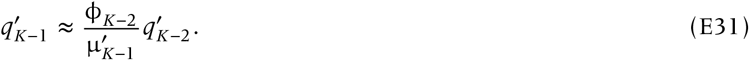

By induction, we have

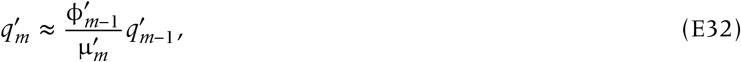

and thus

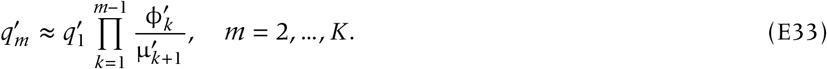

Substituting these expressions into the dynamics of class 1 leads to:

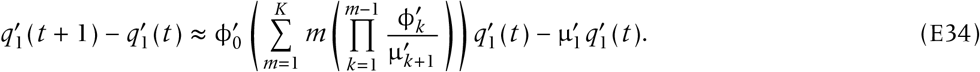

The invasion condition is determined by requiring the right-hand side to be positive, which gives

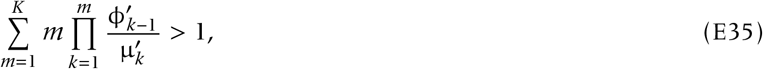

as desired.

##### The Wright-Fisher model and the advantage of the structural approach

The standard approach to evolutionary invasion analysis in subdivided populations, particularly under the Wright-Fisher process with local density dependence, heavily relies on tracking the probability distribution of local patch states (i.e., the number of mutants in a patch). As brilliantly demonstrated by Mullon *et al*. (2016, 2018), one can evaluate the invasion condition by differentiating the lineage fitness. However, because the Wright-Fisher process yields a dense transition matrix, analytically solving the recursions for the perturbed mutant distributions or identical-by-descent (IBD) probabilities requires remarkable mathematical effort and biological intuition. This complexity severely limits the generalizability of the traditional approach when the model incorporates additional class structures such as sex, age, or environmental heterogeneity.

Here, we demonstrate how our structural framework, specifically the synthesis of the projected next-generation matrix (PNGM) and the invasion determinant, effortlessly bypasses this high-dimensional hurdle.

We start by constructing the full *K* × *K* next-generation matrix *G* for a focal patch, where the state *k* ∈ {1, 2, …, *K*} represents the number of mutants in the patch. We biologically decompose the NGM into two distinct pathways: local philopatric transitions *S* and global seeding of new patches *F*, such that

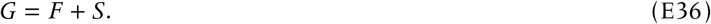

The matrix *S* describes the local demography (e.g., binomial sampling in the Wright-Fisher process) where mutants stay in the focal patch. Importantly, ρ(*S*) < 1 because without global seeding, the local mutant lineage is doomed to ultimate extinction under the infinite-island assumption.

The matrix *F* describes the production of successful emigrants that colonize resident patches. In the infinite-island model, a dispersing mutant will almost surely land in a patch entirely occupied by residents, creating a new local lineage starting with exactly one mutant. This biological feature implies a rank-one structure for *F*:

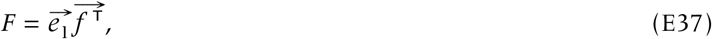

where 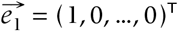 is the unit vector representing the new state of exactly one mutant, and 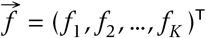 is the vector whose *k*-th element represents the expected number of successful global seedings produced by a patch currently containing *k* mutants.

Under the traditional method, evaluating the spectral radius ρ(*G*) > 1 directly is analytically prohibitive for a dense matrix. However, using the invasion determinant, the invasion condition is equivalent to − det(*I* − *G*) > 0. Substituting the rank-one structure into the determinant and applying the matrix determinant lemma (the Sherman-Morrison formula), we obtain:

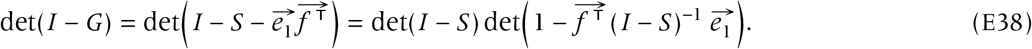

Since ρ(*S*) < 1, we have det(*I* − *S*) > 0. Thus, the invasion condition collapses exactly to a single scalar inequality:

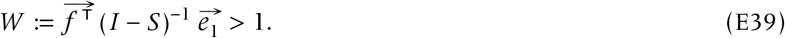

This scalar *W* is precisely the target reproduction number (or type-reproduction number) associated with global seeding. As established in our framework, this metric is invadability-equivalent to the 1 × 1 PNGM projected onto the state of a newly seeded patch (i.e., primary group 𝕏 = {1}), the so-called metapopulation fitness (Metz & Gyllenberg 2001; Ajar 2003; Massol *et al*. 2009; Mullon *et al*. 2016). We have entirely eliminated the need to explicitly track the *K* × *K* mutant frequency dynamics. The matrix (*I* − *S*)^−1^ is the fundamental matrix, where the *k*-th entry of 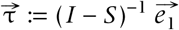 simplyrepresents the expected sojourn time in state *k* starting from a single founder.

The true power of this structural approach shines under weak selection (first-order effects). Instead of solving complex recursions for the perturbed mutant distributions 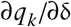 as required in previous studies (Mullon *et al*. 2016, 2018), we can mechanically Taylor-expand the scalar fitness *W*. Let δ be the small phenotypic deviation of the mutant. We expand *S* ≈ *S*^◦^ + δ*S*^(1)^ and 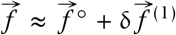. Using the standard matrix identity for the derivative of an inverse, (*I* − *S*)^−1^ ≈ (*I* − *S*^◦^)^−1^ + δ (*I* − *S*^◦^)^−1^ *S*^(1)^ (*I* − *S*^◦^)^−1^, the selection gradient emerges automatically:

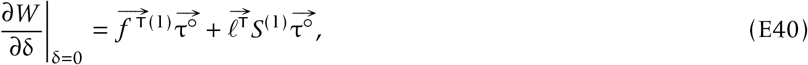

where 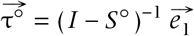 is the neutral sojourn time vector, and 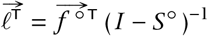 is proportional to (1, 2, …, *K*)^T^.

In this formulation, the term 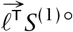 elegantly and automatically generates the inclusive fitness effects (kin selection). Under neutrality, all individuals are selectively equivalent and thus have the exact same individual reproductive value. Consequently, the total reproductive value of a patch (which corresponds to the class state in our NGM) is strictly proportional to the number of mutants it contains, enforcing 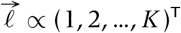. Thus, calculating 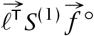 algebraically reduces to evaluating the neutral moments 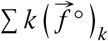 and 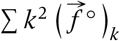, which are fundamentally linked to standard, readily available identity-by-descent (IBD) probabilities or relatedness coefficients.

Under Wright-Fisher demography the local transition matrix is dense, so the convenient tri-diagonal recursions above no longer apply; nevertheless, the metapopulation-fitness criterion still reduces to the scalar condition.

## Notes

### Competing Interest Statement

The authors have declared no competing interest.

### Summary of Updates

Condensed the whole text, removing some parts from the discussion, and changed the figure structures

### Funding

- Japan Society for the Promotion of Science: 24H01528, 24H02291

## References

Abe, J., Iritani, R., Tsuchida, K., Kamimura, Y., & West, S. A. (2021). A solution to a sex ratio puzzle in *Melittobia* wasps. Proc. Natl. Acad. Sci. USA, 118.20, e2024656118. DOI: 10.1073/pnas.2024656118.

Ajar, É. (2003). Analysis of disruptive selection in subdivided populations. BMC Evol. Biol., 3.1, p. 22. DOI: 10.1186/1471-2148-3-22.

Boots, M. & Best, A. (2018). The evolution of constitutive and induced defences to infectious disease. Proc. R. Soc. B, 285.1883. DOI: 10.1098/rspb.2018.0658.

Buckingham, L. J., Bruns, E. L., & Ashby, B. (2023). The evolution of age-specific resistance to infectious disease. Proc. R. Soc. B, 290.1991. DOI: 10.1098/rspb.2022.2000.

Caswell, H. (2001). Matrix population models. Wiley Online Library.

Charnov, E. L. (1976). Optimal foraging, the marginal value theorem. Theor. Popul. Biol., 9.2, pp. 129–136. DOI: 10.1016/0040-5809(76)90040-x.

Christiansen, F. B. (1991). On conditions for evolutionary stability for a continuously varying character. Am. Nat., pp. 37–50. DOI: 10.1086/285203.

Coulson, T., Benton, T., Lundberg, P., Dall, S., Kendall, B., & Gaillard, J.-M. (2005). Estimating individual contributions to population growth: evolutionary fitness in ecological time. Proc. R. Soc. B, 273.1586, pp. 547–55. DOI: 10.1098/rspb.2005.3357.

Coulson, T., MacNulty, D. R., Stahler, D. R., vonHoldt, B., Wayne, R. K., & Smith, D. W. (2011). Modeling effects of environmental change on wolf population dynamics, trait evolution, and life history. Science, 334.6060, pp. 1275–1278. DOI: 10.1126/science.1209441.

Coulson, T., Tuljapurkar, S., & Childs, D. Z. (2010). Using evolutionary demography to link life history theory, quantitative genetics and population ecology. J. Anim. Ecol., 79.6, pp. 1226–1240. DOI: 10.1111/j.1365-2656.2010.01734.x.

Courteau, J. & Lessard, S. (2000). Optimal sex ratios in structured populations. J. Theor. Biol., 207.2, pp. 159–175. DOI: 10.1006/jtbi.2000.2160.

Cushing, J. M. (1998). An Introduction to Structured Population Dynamics. Society for Industrial and Applied Mathematics. DOI: 10.1137/1.9781611970005.

Day, T. & Burns, J. G. (2003). A consideration of patterns of virulence arising from host-parasite coevolution. Evolution, 57.3, pp. 671–676. DOI: 10.1111/j.0014-3820.2003.tb01558.x.

de Camino Beck, T. & Lewis, M. A. (2007). A new method for calculating net reproductive rate from graph reduction with applications to the control of invasive species. Bull. Math. Biol., 69.4, pp. 1341–1354. DOI: 10.1007/s11538-006-9162-0.

Dieckmann, U. & Law, R. (1996). The dynamical theory of coevolution: a derivation from stochastic ecological processes. J. Math. Biol., 34.5-6, pp. 579–612. DOI: 10.1007/s002850050022.

Diekmann, O., Heesterbeek, J. A. P., & Metz, J. A. J. (1990). On the definition and the computation of the basic reproduction ratio *R*0 in models for infectious diseases in heterogeneous populations. J. Math. Biol., 28.4. DOI: 10.1007/bf00178324.

Eshel, I. (1996). On the changing concept of evolutionary population stability as a reflection of a changing point of view in the quantitative theory of evolution. J. Math. Biol., 34.5–6, pp. 485–510. DOI: 10.1007/bf02409747.

Ewens, W. J. (2004). Mathematical Population Genetics: I. Theoretical Introduction. 2nd ed. Interdisciplinary Applied Mathematics 27. Springer-Verlag New York.

Fisher, R. A. (1930). The genetical theory of natural selection. Clarendon Press. DOI: 10.5962/bhl.title.27468.

Fisher, R. A. (1941). Average excess and average effect of a gene substitution. Ann. Eugen., 11.1, pp. 53–63. DOI: 10.1111/j.1469-1809.1941.tb02272.x.

Frank, S. A. (1998). Foundations of social evolution. Princeton University Press.

Geritz, S. A. H., Kisdi, É., Meszéna, G., & Metz, J. (1998). Evolutionarily singular strategies and the adaptive growth and branching of the evolutionary tree. Evol. Ecol., 12.1, pp. 35–57. DOI: 10.1023/a:1006554906681.

Geritz, S. A. H. (2005). Resident-invader dynamics and the coexistence of similar strategies. J. Math. Biol., 50.1, pp. 67–82. DOI: 10.1007/s00285-004-0280-8.

Geritz, S. A. H., Gyllenberg, M., Jacobs, F. J., & Parvinen, K. (2002). Invasion dynamics and attractor inheritance. J. Math. Biol., 44.6, pp. 548–560. DOI: 10.1007/s002850100136.

Giaimo, S. (2025). Elasticity analysis of population growth: Implications of matrix model construction. Ecol. Model., 507, p. 111163. DOI: 10.1016/j.ecolmodel.2025.111163.

Giaimo, S., Arranz, J., & Traulsen, A. (2018). Invasion and effective size of graph-structured populations. PLOS Comput. Biol., 14.11, e1006559. DOI: 10.1371/journal.pcbi.1006559.

Giaimo, S. & Traulsen, A. (2019). Generation Time Measures the Trade-Off between Survival and Reproduction in a Life Cycle. Am. Nat., 194.2, pp. 285–290. DOI: 10.1086/704155.

Giaimo, S. & Traulsen, A. (2021). Applying symmetries of elasticities in matrix population models. Theor. Ecol., 14.3, pp. 359–366. DOI: 10.1007/s12080-021-00513-x.

Hamilton, W. D. & May, R. M. (1977). Dispersal in stable habitats. Nature, 269.5629, pp. 578–581. DOI: 10.1038/269578a0.

Hastings, A. & Botsford, L. W. (2006a). A simple persistence condition for structured populations. Ecol. Lett., 9.7, pp. 846–852. DOI: 10.1111/j.1461-0248.2006.00940.x.

Hastings, A. & Botsford, L. W. (2006b). Persistence of spatial populations depends on returning home. Proc. Natl. Acad. Sci. USA, 103.15, pp. 6067–6072. DOI: 10.1073/pnas.0506651103.

Hofbauer, J. & Sigmund, K. (1990). Adaptive dynamics and evolutionary stability. Appl. Math. Lett., 3.4, pp. 75–79. DOI: 10.1016/0893-9659(90)90051-c.

Hurford, A., Cownden, D., & Day, T. (2010). Next-generation tools for evolutionary invasion analyses. J. R. Soc. Interface, 7.45, pp. 561–571. DOI: 10.1098/rsif.2009.0448.

Inaba, H. & Nishiura, H. (2008). The state-reproduction number for a multistate class age structured epidemic system and its application to the asymptomatic transmission model. Math. Biosci., 216.1, pp. 77–89. DOI: 10.1016/j.mbs.2008.08.005.

Iritani, R. & Cheptou, P.-O. (2017). Joint evolution of differential seed dispersal and self-fertilization. J. Evol. Biol., 30.8, pp. 1526–1543. DOI: 10.1111/jeb.13120.

Iritani, R. (2020). Gametophytic competition games among relatives: when does spatial structure select for facilitativeness or competitiveness in pollination? J. Ecol., 108.1, pp. 1–13. DOI: 10.1111/1365-2745.13282.

Iritani, R. & Iwasa, Y. (2014). Parasite infection drives the evolution of state-dependent dispersal of the host. Theor. Popul. Biol., 92, pp. 1–13. DOI: 10.1016/j.tpb.2013.10.005.

Iritani, R., Visher, E., & Boots, M. (2019). The evolution of stage-specific virulence: Differential selection of parasites in juveniles. Evol. Lett., 3.2, pp. 162–172. DOI: 10.1002/evl3.105.

Iritani, R., West, S. A., & Abe, J. (2021). Cooperative interactions among females can lead to even more extraordinary sex ratios. Evol. Lett., 5.4, pp. 370–384. DOI: 10.1002/evl3.217.

Jones, O. R., Barks, P., Stott, I., James, T. D., Levin, S., Petry, W. K., Capdevila, P., CheCastaldo, J., Jackson, J., Römer, G., Schuette, C., Thomas, C. C., & SalgueroGómez, R. (2022). Rcompadre and Rage—Two R packages to facilitate the use of the COMPADRE and COMADRE databases and calculation of lifehistory traits from matrix population models. Methods Ecol. Evol., 13.4, pp. 770–781. DOI: 10.1111/2041-210x.13792.

Kemeny, J. G., Snell, J., & Knapp, A. (1966). Markov chains. New York, NY: University Series in Higher Mathematics, Van Nostrand.

Kruuk, L. & Hill, W. (2008). Evolutionary dynamics of wild populations: the use of long-term pedigree data. Proc. R. Soc. B, 275.1635, pp. 593–596. DOI: 10.1098/rspb.2007.1689.

Kuijper, B. & Johnstone, R. A. (2019). The evolution of early-life effects on social behaviour—why should social adversity carry over to the future? Phil. Trans. R. Soc. B, 374.1770, p. 20180111. DOI: 10.1098/rstb.2018.0111.

Lefkovitch, L. P. (1965). The study of population growth in organisms grouped by stages. Biometrics, 21.1, p. 1. DOI: 10.2307/2528348.

Lewis, M. A., Shuai, Z., & van den Driessche, P. (2019). A general theory for target reproduction numbers with applications to ecology and epidemiology. J. Math. Biol., DOI: 10.1007/s00285-019-01345-4.

Li, C.-K. & Schneider, H. (2002). Applications of Perron-Frobenius theory to population dynamics. J. Math. Biol., 44.5, pp. 450–462. DOI: 10.1007/s002850100132.

Lion, S. (2018). Class structure, demography, and selection: reproductive-value weighting in nonequilibrium, polymorphic populations. Am. Nat., 191.5, pp. 620–637. DOI: 10.1086/696976.

Lion, S., Boots, M., & Sasaki, A. (2022). Multi-morph eco-evolutionary dynamics in structured populations. Am. Nat., 200.3, pp. 345–372. DOI: 10.1086/720439.

Lion, S. & Gandon, S. (2022). Evolution of class-structured populations in periodic environments. Evolution, 76.8, pp. 1674–1688. DOI: 10.1111/evo.14522.

Lion, S. & Metz, J. A. J. J. (2018). Beyond *R*0 maximisation: on pathogen evolution and environmental dimensions. Trends Ecol. Evol., 33, pp. 75–90. DOI: 10.1016/j.tree.2018.02.004.

Lion, S., Sasaki, A., & Boots, M. (2023). Extending ecoevolutionary theory with oligomorphic dynamics. Ecol. Lett., 26.S1. DOI: 10.1111/ele.14183.

Massol, F., Calcagno, V., & Massol, J. (2009). The metapopulation fitness criterion: Proof and perspectives. Theor. Popul. Biol., 75.2–3, pp. 183–200. DOI: 10.1016/j.tpb.2009.02.005.

Massol, F. & Cheptou, P.-O. (2011). Evolutionary syndromes linking dispersal and mating system: the effect of autocorrelation in pollination conditions. Evolution, 65.2, pp. 591–598. DOI: 10.1111/j.1558-5646.2010.01134.x.

Massol, F. & Débarre, F. (2015). Evolution of dispersal in spatially and temporally variable environments: The importance of life cycles. Evolution, 69.7, pp. 1925–1937. DOI: 10.1111/evo.12699.

Maynard Smith, J. (1982). Evolution and the Theory of Games. Cambridge university press.

Maynard Smith, J. & Price, G. R. (1973). The logic of animal conflict. Nature, 246, p. 15. DOI: 10.1038/246015a0.

Metz, J. A. J. & Leimar, O. (2011). A simple fitness proxy for structured populations with continuous traits, with case studies on the evolution of haplo-diploids and genetic dimorphisms. J. Biol. Dynam., 5.2, pp. 163–190. DOI: 10.1080/17513758.2010.502256.

Metz, J. A. & Gyllenberg, M. (2001). How should we define fitness in structured metapopulation models? Including an application to the calculation of evolutionarily stable dispersal strategies. Proc. R. Soc. B, 268.1466, pp. 499–508.

Metz, J. A., Nisbet, R., & Geritz, S. (1992). How should we define ‘fitness’ for general ecological scenarios? Trends Ecol. Evol., 7.6, pp. 198–202.

Mitchell, E., Graham, A. L., Úbeda, F., & Wild, G. (2022). On maternity and the stronger immune response in women. Nat. Commun., 13.1. DOI: 10.1038/s41467-022-32569-6.

Morita, K., Sasaki, A., & Iritani, R. (2025). How can interspecific pollen transfer affect the coevolution and coexistence of two closely related plant species? Oikos, e11133. DOI: 10.1002/oik.11133.

Mullon, C., Keller, L., & Lehmann, L. (2016). Evolutionary stability of jointly evolving traits in subdivided populations. Am. Nat., 188.2, pp. 175–195. DOI: 10.1086/686900.

Mullon, C. & Lehmann, L. (2014). The robustness of the weak selection approximation for the evolution of altruism against strong selection. J. Evol. Biol., 27.10, pp. 2272–2282. DOI: 10.1111/jeb.12462.

Pelletier, F., Clutton-Brock, T., Pemberton, J., Tuljapurkar, S., & Coulson, T. (2007). The Evolutionary Demography of Ecological Change: Linking Trait Variation and Population Growth. Science, 315.5818, pp. 1571–1574. DOI: 10.1126/science.1139024.

Priklopil, T. & Lehmann, L. (2020). Invasion implies substitution in ecological communities with class-structured populations. Theor. Popul. Biol., 134, pp. 36–52. DOI: 10.1016/j.tpb.2020.04.004.

Priklopil, T. & Lehmann, L. (2024). On the interpretation of the operation of natural selection in class-structured populations. Am. Nat., 203.2, pp. 292–304. DOI: 10.1086/727970.

Rees, M. & Ellner, S. P. (2016). Evolving integral projection models: evolutionary demography meets eco-evolutionary dynamics. Methods Ecol. Evol., 7.2, pp. 157–170. DOI: 10.1111/2041-210x.12487.

Roberts, M. G. & Heesterbeek, J. A. P. (2003). A new method for estimating the effort required to control an infectious disease. Proc. R. Soc. B, 270.1522, pp. 1359–1364. DOI: 10.1098/rspb.2003.2339.

Rodrigues, A. M. M. & Gardner, A. (2015). Simultaneous failure of two sex-allocation invariants: implications for sex-ratio variation within and between populations. Proc. R. Soc. B, 282.1810, p. 20150570. DOI: 10.1098/rspb.2015.0570.

Rodrigues, A. M. M. & Gardner, A. (2013). Evolution of helping and harming in heterogeneous groups. Evolution, 67.8, pp. 2284–2298. DOI: 10.1111/j.1558-5646.2012.01594.x.

Rousset, F. (2004). Genetic Structure and Selection in Subdivided Populations (MPB-40). Princeton University Press.

Rueffler, C. & Metz, J. A. J. (2013). Necessary and sufficient conditions for *R*0 to be a sum of contributions of fertility loops. J. Math. Biol., 66.4-5, pp. 1099–1122. DOI: 10.1007/s00285-012-0575-0.

Rueffler, C., Metz, J. A. J., & Van Dooren, T. J. M. (2012). What life cycle graphs can tell about the evolution of life histories. J. Math. Biol., 66.1-2, pp. 225–279. DOI: 10.1007/s00285-012-0509-x.

Salguero-Gómez, R., Jones, O. R., Archer, C. R., Bein, C., Buhr, H. de, Farack, C., Gottschalk, F., Hartmann, A., Henning, A., Hoppe, G., Römer, G., Ruoff, T., Sommer, V., Wille, J., Voigt, J., Zeh, S., Vieregg, D., Buckley, Y. M., Che□Castaldo, J., Hodgson, D., Scheuerlein, A., Caswell, H., & Vaupel, J. W. (2016). COMADRE □: a global data base of animal demography. J. Anim. Ecol., 85.2, pp. 371–384. DOI: 10.1111/1365-2656.12482.

Salguero-Gómez, R., Jones, O. R., Archer, C. R., Buckley, Y. M., Che-Castaldo, J., Caswell, H., Hodgson, D., Scheuerlein, A., Conde, D. A., Brinks, E., Buhr, H. de, Farack, C., Gottschalk, F., Hartmann, A., Henning, A., Hoppe, G., Römer, G., Runge, J., Ruoff, T., Wille, J., Zeh, S., Davison, R., Vieregg, D., Baudisch, A., Altwegg, R., Colchero, F., Dong, M., Kroon, H. de, Lebreton, J.-D., Metcalf, C. J. E., Neel, M. M., Parker, I. M., Takada, T., Valverde, T., Vélez-Espino, L. A., Wardle, G. M., Franco, M., & Vaupel, J. W. (2014). The COMPADRE Plant Matrix Database: an open online repository for plant demography. J. Ecol., 103.1, pp. 202–218. DOI: 10.1111/1365-2745.12334.

Sasaki, A. & Dieckmann, U. (2011). Oligomorphic dynamics for analyzing the quantitative genetics of adaptive speciation. J. Math. Biol., 63.4, pp. 601–635. DOI: 10.1007/s00285-010-0380-6.

Shefferson, R. P. (2025). adapt3: Adaptive Dynamics and Community Matrix Model Projections. DOI: 10.32614/cran.package.adapt3.

Sigmund, K. (2010). The calculus of selfishness. Princeton University Press.

Smallegange, I. M. & Coulson, T. (2013). Towards a general, population-level understanding of eco-evolutionary change. Trends Ecol. Evol., 28.3, pp. 143–148. DOI: 10.1016/j.tree.2012.07.021.

Stott, I., Townley, S., Carslake, D., & Hodgson, D. J. (2010). On reducibility and ergodicity of population projection matrix models. Methods Ecol. Evol., 1.3, pp. 242–252. DOI: 10.1111/j.2041-210x.2010.00032.x.

Takada, T. & Kigami, J. (1991). The dynamical attainability of ESS in evolutionary games. J. Math. Biol., 29.6, pp. 513–529. DOI: 10.1007/bf00164049.

Takada, T. & Nakajima, H. (1992). An analysis of life history evolution in terms of the density-dependent Lefkovitch matrix model. Math. Biosci., 112.1, pp. 155–176. DOI: 10.1016/0025-5564(92)90091-a.

Taylor, P. D. (1990). Allele-frequency change in a class-structured population. Am. Nat., 135, pp. 95–106. DOI: 10.1086/285034.

Taylor, P. D. & Bulmer, M. G. (1980). Local mate competition and the sex ratio. J. Theor. Biol., 86.3, pp. 409–419. DOI: 10.1016/0022-5193(80)90342-2.

Taylor, P. D. & Frank, S. A. (1996). How to make a kin selection model. J. Theor. Biol., 180.1, pp. 27–37. DOI: 10.1006/jtbi.1996.0075.

Úbeda, F. & Jansen, V. A. A. (2016). The evolution of sex-specific virulence in infectious diseases. Nat. Commun., 7, p. 13849. DOI: 10.1038/ncomms13849.

van den Driessche, P. & Watmough, J. (2002). Reproduction numbers and sub-threshold endemic equilibria for compartmental models of disease transmission. Math. Biosci., 180.1, pp. 29–48. DOI: 10.1016/s0025-5564(02)00108-6.

Wickman, J., Koffel, T., & Klausmeier, C. A. (2023). A theoretical framework for trait-based eco-evolutionary dynamics: population structure, intraspecific variation, and community assembly. Am. Nat., 201.4, pp. 501–522. DOI: 10.1086/723406.

Williams, P. D. & Kamel, S. J. (2021). Evolutionary invasion analysis in structured populations. Evol. Biol., 48.4, pp. 422–427. DOI: 10.1007/s11692-021-09547-9.

Wright, S. (1931). Evolution in Mendelian populations. Genetics, 16.2, p. 97. DOI: 10.1007/bf02459575.

Zurita-Gutiérrez, Y. H. & Lion, S. (2015). Spatial structure, host heterogeneity and parasite virulence: implications for vaccine-driven evolution. Ecol. Lett., 18.8, pp. 779–789. DOI: 10.1111/ele.12455.

## References

Avila, P. & Mullon, C. (2023). Evolutionary game theory and the adaptive dynamics approach: adaptation where individuals interact. Phil. Trans. R. Soc. B, 378.1876. DOI: 10.1098/rstb.2021.0502.

Berman, A. & Plemmons, R. J. (1987). Nonnegative matrices in the mathematical sciences. Classics in applied mathematics 9. Society for Industrial and Applied Mathematics.

Crabtree, D. E. & Haynsworth, E. V. (1969). An identity for the Schur complement of a matrix. Proc. Am. Math. Soc., 22.2, pp. 364–366. DOI: 10.1090/s0002-9939-1969-0255573-1.

Frank, S. A. (1998). Foundations of social evolution. Princeton University Press.

Graham, A. (2018). Kronecker products and matrix calculus with applications. Courier Dover Publications.

Horn, R. A. & Johnson, C. R. (1985). Matrix Analysis. 169. Cambridge University Press, Cambridge.

Mullon, C., Keller, L., & Lehmann, L. (2018). Social polymorphism is favoured by the co-evolution of dispersal with social behaviour. Nat. Ecol. Evol., 2.1, p. 132. DOI: 10.1038/s41559-017-0397-y.

Varga, R. S. (1963). Matrix iterative analysis. Graduate Texts in Mathematics. Prentice–Hall, Englewood Cliffs, NJ.

